# Organ-restricted vascular delivery of nanoparticles for lung cancer therapy

**DOI:** 10.1101/2020.03.05.969212

**Authors:** Deniz A. Bölükbas, Stefan Datz, Charlotte Meyer-Schwickerath, Carmela Morrone, Ali Doryab, Dorothee Gößl, Malamati Vreka, Lin Yang, Christian Argyo, Sabine H. van Rijt, Michael Lindner, Oliver Eickelberg, Tobias Stoeger, Otmar Schmid, Sandra Lindstedt, Georgios T. Stathopoulos, Thomas Bein, Darcy E. Wagner, Silke Meiners

## Abstract

Nanomedicines hold immense promise for a number of devastating diseases due to the ability to custom-design both the carrier and cargo. However, their clinical implementation has been hampered by physicochemical and biological barriers and off-target deposition which impair cell specific targeting, especially in internal organs. This study reports a new delivery approach using organ-restricted vascular delivery to allow for direct administration and recirculation of stimuli-responsive nanoparticles to promote cellular uptake into an organ of interest. Using this technique, nanoparticles reach the interior of dense tumors and are selectively taken up by lung cancer cells. Importantly, this surgical approach is essential as the same nanoparticles do not reach lung tumor cells upon systemic or intratracheal administration. Organ-restricted vascular delivery thus opens up new avenues for optimized nanotherapies for cancer and other diseases.

## 1. Introduction

The use of nanoparticles as therapeutic agents for cancer therapy and other diseases has attracted major attention in the past decades. The first generation of nanomedicines for cancer relied on passive targeting of cancer cells based on the concept of enhanced permeability and retention (EPR) effect observed in tumors.^[1]^ The EPR effect has been described to promote preferential accumulation of nanoparticles in the tumor due to increased blood vessel permeability and impaired lymphatic drainage in the tumorous regions. Several FDA-approved nanomedicines have been designed with the intent of exerting their therapeutic effect by passive targeting of tumors.^[2]^ More recently, nanoparticles incorporating active targeting strategies have emerged as an alternative approach to fine-tune nanoparticle delivery to specific cell types.^[3]^ To achieve cell-specific targeting, nanoparticles can be functionalized with targeting agents on their surface, which can bind to receptors specifically overexpressed on tumor or tumor-associated cells. Furthermore, smart nanoparticles which can respond to specific external stimuli and release of cargo into the tumor have been explored to limit off-target effects.^[4]^ However, the initial excitement accompanying these innovative nanomedical approaches has failed to translate into clinical success, largely due to physicochemical and biological factors which impair nanoparticle targeting *in vivo.*^[5]^

In particular, increased interstitial hypertension in tumors is known to reduce convective transport and the dense extracellular matrix of the tumor environment hinders the diffusion of particles.^[6]^ Furthermore, nanoparticles can be directly engulfed or immunologically tagged for destruction by the mononuclear phagocyte system (MPS) to be cleared in the liver or spleen, or by alveolar macrophages in the lung.^[7]^ Accordingly, up to 99% of all systemically administered nanoparticles are deposited in the liver and spleen, representing a major limiting factor for effective application of nanomedicines.^[8]^ In order to increase nanoparticle accumulation in solid tumors, clearance of particles by competing organs needs to be minimized and novel organ-specific methods of nanoparticle delivery should be considered.^[9]^ While local delivery is possible for tumors of peripheral organs, such as the skin, delivery into tumors of internal organs, such as the lung, remains challenging.

Among solid tumors, nanoparticle delivery for lung cancer is one of the most challenging with the lowest targeting efficacy compared to other solid cancer types.^[8]^ Lung cancer affects over 1.7 million patients per year and has the highest mortality rate among all cancers worldwide.^[10]^ Although recent advances in cancer immunotherapy ^[11]^ and targeted therapy ^[12]^ have resulted in encouraging progress, the number of patients and mortalities is expected to rise for lung cancer, mainly due to increases in smoking rates and exposure to pollution. Previously, there have been attempts to locally deliver chemotherapeutics to lung tumors through either the airways ^[13]^ or the vasculature.^[14]^ While airway delivery via inhalation has shown some degree of efficacy, particle delivery into the core of larger, solid tumors has not yet been achieved and thus inhalation therapies are limited to the treatment of smaller tumors. On the other hand, vascular delivery of chemotherapeutics exclusively to the lung, termed as isolated lung perfusion, is feasible in patients with lung tumors as demonstrated previously.^[15]^ However, these approaches are infrequent due to high levels of pulmonary toxicity resulting from chemotherapeutics accumulating not only in tumors but also in healthy lung tissue.^[14]^

Nanotherapy is particularly well suited for overcoming pulmonary toxicity associated with healthy tissue since with the help of the EPR effect and the recently described transcytotic pathway upregulated in endothelial cells lining the tumor vasculature ^[16]^, nanoparticles could be successfully and selectively delivered to tumorous regions. Here, we provide proof-of-concept evidence for effective delivery of both active and passive targeting nanoparticles with pH-responsiveness into the interior of solid, dense lung tumors using a novel organ-restricted vascular delivery (ORVD) strategy. This approach overcomes unwanted accumulation of nanoparticles in off-target organs such as the liver and spleen and in alveolar macrophages of the lung by nanoparticle delivery into the surgically isolated vasculature of the organ of interest. It also eliminates extravasation of chemotherapeutics into healthy regions of the lung which is associated with pulmonary toxicity after isolated lung perfusion (**Figure 1**). Furthermore, ORVD results in cellular uptake and endosomal escape of nanoparticles in lung tumor cells, which is a prerequisite for nanoparticle-mediated drug delivery. ORVD is a versatile new delivery strategy which overcomes physicochemical and biological barriers for local delivery of nanomedicines to internal organs.

**Figure 1.**
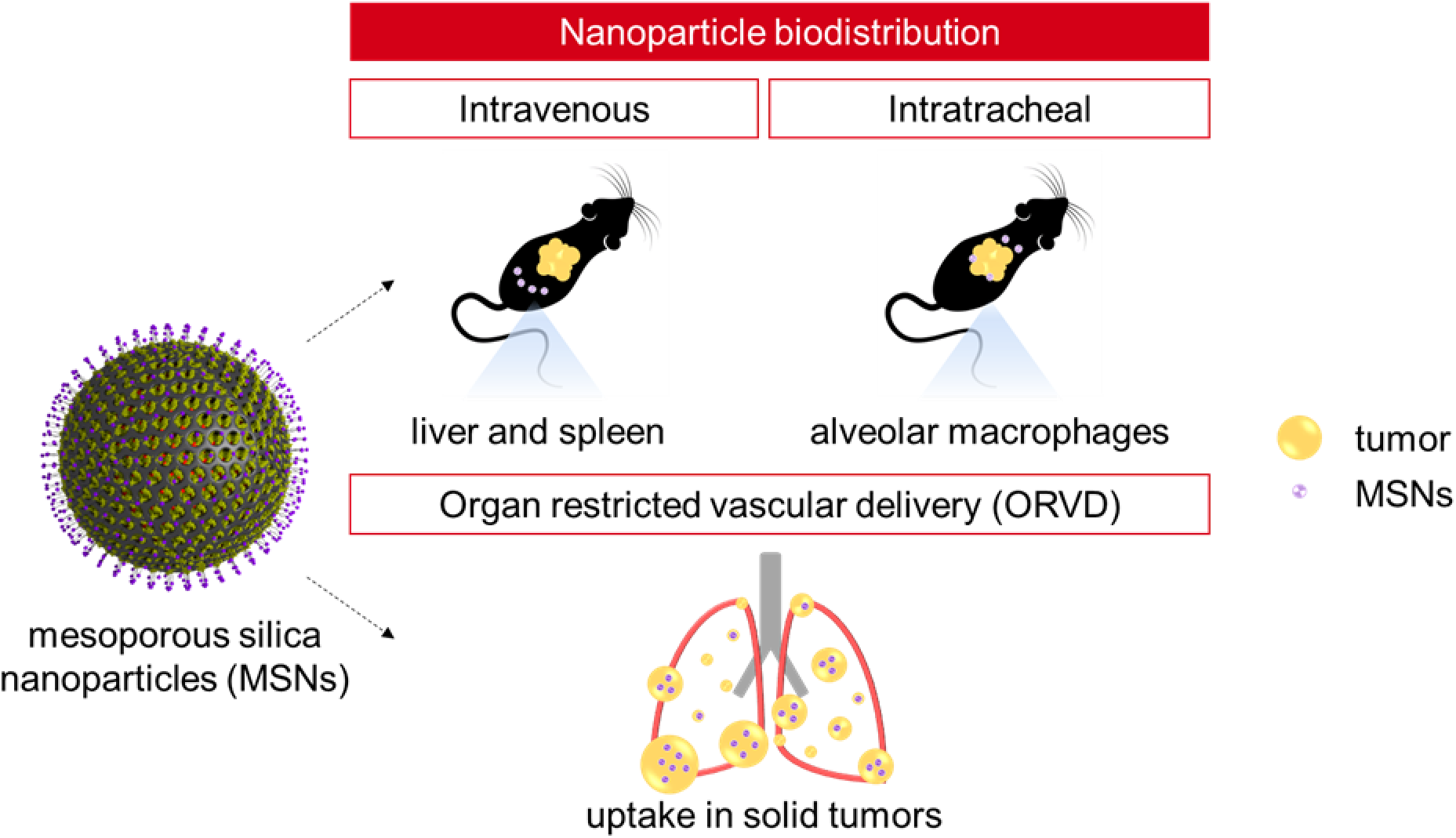
Experimental overview comparing different delivery routes for application of mesoporous silica nanoparticles (MSNs) for lung cancer therapy. Conventional delivery methods (*i.e.,* intravenous and intratracheal) present barriers for effective delivery of nanoparticles for lung cancer. Implementation of organ-restricted vascular delivery (ORVD) of MSNs as a novel approach to overcome the physicochemical and biological barriers with isolated nanoparticle recirculation at a controlled and effective flow rate, synergizing the enhanced permeability and retention (EPR) effect with the absence of the mononuclear phagocyte system (MPS).

## 2. Results

### 2.1. Synthesis and characterization of mesoporous silica nanoparticles

We first generated functionalized mesoporous silica nanoparticles (MSNs) ^[17]^ which can be used for active or passive targeting of specific tumor cells. Additionally, our MSNs contain a pH-responsive element allowing selective release into the tumor microenvironment or upon internalization into the endolysosomal compartment. The internal pore system of the MSNs was functionalized with thiol groups and the external particle surface with amino groups, creating a nanoparticle platform which allows for a wide range of customizable functionalizations (**Figure 2A**). The pH-cleavable linker system, containing a biotin functionality, was added to the external surface of the MSN. The glycoprotein avidin was attached to the outer surface of the MSN_Biotin_ via noncovalent association with the biotin groups (MSN_AVI_), acting as a stable gatekeeper of the pore system. MSN_AVI_ nanoparticles without further functionalization were used for passive targeting of tumors in this study. Different targeting ligands can be attached to the outer surface of the avidin gatekeepers for active targeting of tumors (Figure 2B and Figure S1A).

**Figure 2.**
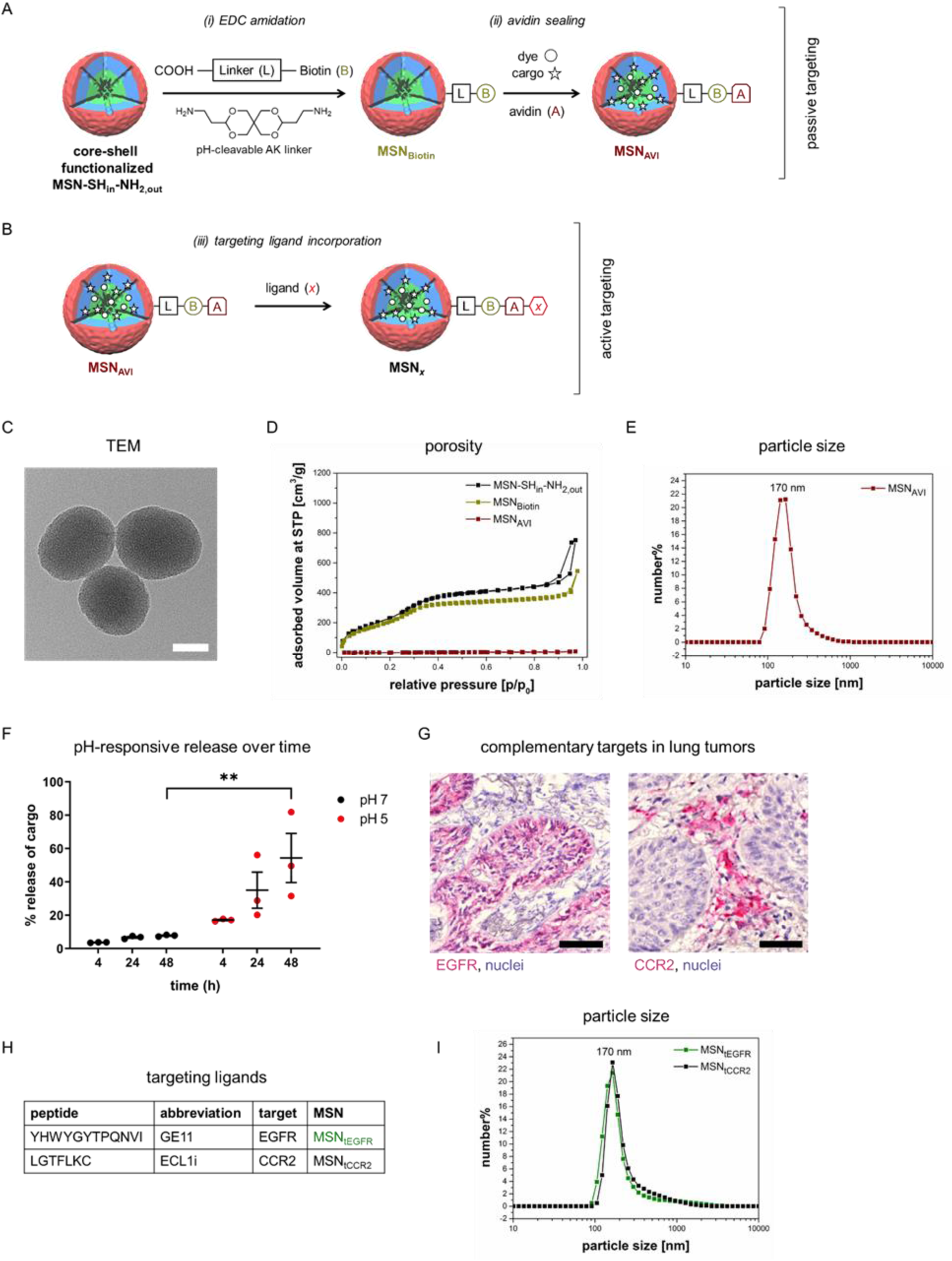
Synthesis and characterization of mesoporous silica nanoparticles for passive and active targeting. a) Delayed co-condensation process leads to different core (green, thiol groups) and shell (red, amino groups) functionalization of MSN-SH_in_-NH_2,out_. (i) MSN_Biotin_ nanoparticles were generated by first transforming amino groups into carboxy groups, followed by EDC amidation with the pH-cleavable AK linker, and subsequent addition of covalently bound biotin. (ii) MSN_AVI_ nanoparticles were generated by the addition of avidin to efficiently seal the mesopores after cargo loading and covalent attachment of the dyes at the thiol groups in the inner pore system. b) Functionalized nanoparticles (MSN*_x_*) with active targeting were generated by covalent attachment of specific ligands to the outer surface of MSN_AVI_ for lung cancer. c) Transmission electron micrograph (TEM) of MSN-SH_in_-NH_2,out_. Scale bar = 50 nm. d) Nitrogen sorption isotherms of MSN-SH_in_-NH_2,out_ (black), MSN_Biotin_ (green) and MSN_AVI_ (red) showing efficient sealing of the pores. e) Dynamic light scattering (DLS) of MSN_AVI_ (red) in water showing size uniformity. f) Time-dependent pH-responsive release at pH 7 and pH 5. ** p = 0.0026, two-way ANOVA, Sidak’s multiple comparisons test comparing cargo release between pH 5 and pH 7 over time. Values given are an average of three independent experiments ± standard error of the mean g) Immunohistochemical staining representing complementary distribution of EGFR and CCR2 in a human non-small cell lung cancer (NCSLC) specimen. Scale bar = 50 μm. h) Peptide sequences for EGFR and CCR2 targeting ligands, *i.e.,* GE11 and ECL1i, respectively. i) Dynamic light scattering (DLS) of MSN_tEGFR_ (green) and MSN_tCCR2_ (black) in water showing size stability following functionalization.

Synthesized core-shell functionalized MSNs (MSN-SH_in_-NH_2,out_) were amorphous (Figure S1B) and spherical in shape with an approximate size of around 100-150 nm (Figure 2C). The mesoporous structure was confirmed using nitrogen sorption measurements (Figure 2D), with pore sizes of around 4 nm (Figure S1C). Following internal and external functionalization (internal thiol external amine functionalization for MSN-SH_in_-NH_2,out_, external carboxy functionalization for MSN_COOH_, external pH-responsive AK linker functionalization for MSN_AK_, and external biotin functionalization for MSN_Biotin_) (Figure S1D), pore size and volume changed negligibly (Figure S1E and F). As anticipated, successful avidin capping resulted in a loss of surface porosity concomitant with a decrease in measured surface area, as the internal surfaces were no longer accessible following capping (Figure S1C and G). We also observed a slight increase in the isoelectric point (Figure S1H) and stabilization of surface charge across different pHs (Figure S1I) following avidin capping (Figure S1J). MSN_AVI_ nanoparticles had a mean particle size of 170 nm in aqueous media (Figure 2E), demonstrating colloidal stability. We next sought to confirm functionality of our pH-cleavable linker system using a neutral pH of 7 and a pH of 5, representing the acidic pH of the tumor microenvironment. Propidium iodide (PI) was loaded into MSN_AVI_ (0.365 mg PI/mg MSN_AVI_) and the release of PI was measured over time after the pH was changed from 7 to 5. While we observed minimal PI release under neutral pH over 48 h, acidic pH induced cargo release over time with up to ∼60% release after 48 h (Figure 2F). Release at 48 h at acidic pH was significantly increased as compared to the same time point at neutral pH, indicating the specificity and stability of our pH-cleavable linkage system and avidin capping. Next, we further functionalized our MSN_AVI_ with targeting ligands for two different receptors that are well known to be specifically elevated in distinct compartments of human lung tumors, i.e., epidermal growth factor receptor (EGFR) on lung tumor and tumor-associated cells ^[18]^ or C-C chemokine receptor type 2 (CCR2) on tumor-associated macrophages in the surrounding stroma ^[19]^ (Figure 2G and lower magnification image in Figure S1K). Active targeting of these two receptors by exploiting two different particle types can be regarded as a complementary approach to achieve an additive anti-tumor effect on cancer as well as cancer-associated cells. Here, we functionalized the artificial peptides GE11 ^[20]^ and ECL1i ^[21]^ on MSN_AVI_ to generate particles targeting EGFR (MSN_tEGFR_) and CCR2 (MSN_tCCR2_), respectively (Figure 2H). Importantly, colloidal stability was retained in aqueous solutions following the addition of targeting ligands for both MSN_tEGFR_ and MSN_tCCR2_ (Figure 2I and Figure S2L).

### 2.2. Increased uptake of actively targeted MSNs in vitro

To assess whether our active targeting strategy enhances nanoparticle uptake *in vitro*, we exposed A549 cells, a human lung cancer cell line with high EGFR expression (Figure S2A), to receptor-targeted MSN_tEGFR_ and untargeted MSN_AVI_. We observed significantly enhanced uptake of MSN_tEGFR_ in A549 cells as quantified by confocal microscopy (**Figure 3A**) and flow cytometry (Figure 3B). MSN_tEGFR_ co-localized with EGFR, indicating EGFR-mediated uptake. MSN_tCCR2_ specificity was tested in an alveolar macrophage line, MH-S cells, which have high expression of CCR2 (Figure S2B). We again observed significantly increased uptake of MSN_tCCR2_ in MH-S cells, as measured by confocal microscopy (Figure S2C) and flow cytometry (Figure S2D). In order to further confirm cellular uptake, we performed transmission electron microscopy (TEM) of A549 cells exposed to MSN_tEGFR_. Interestingly, we observed both receptor-mediated and unspecific endocytic uptake as well as endosomal escape and evidence of particle degradation (Figure 3C). Together, these results show that the enhanced cellular uptake of the actively targeted nanoparticles *in vitro* is at least partially mediated via receptor-mediated uptake and confirms the functionality of our active targeting strategy.

**Figure 3.**
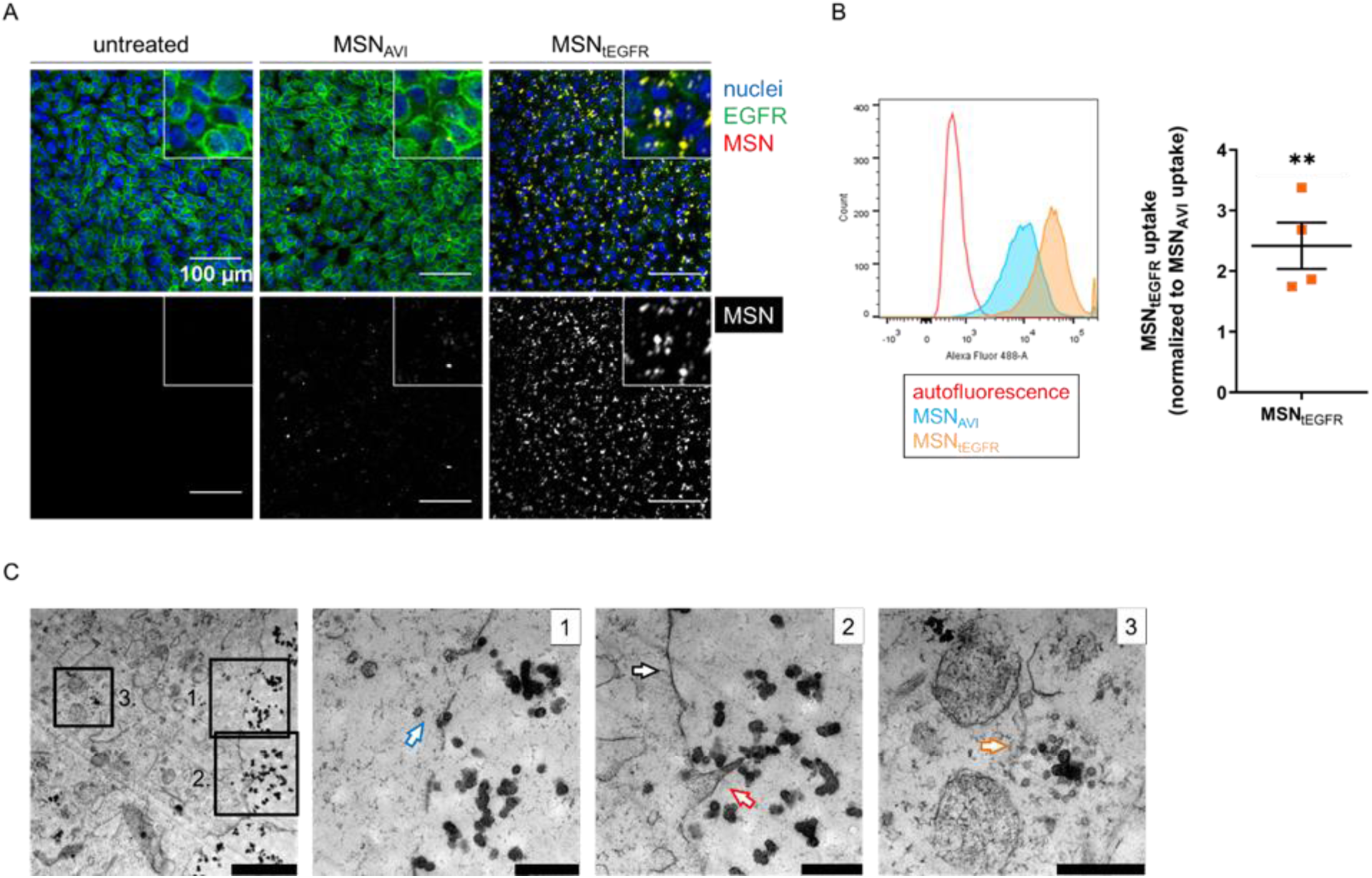
Increased uptake of EGFR targeted MSNs in human lung cancer cells *in vitro*. a) Untargeted *versus* EGFR-targeted uptake of ATTO 633-labeled MSN_AVI_ and MSN_tEGFR_ (red in the upper panel, gray in the lower panel) by human lung cancer cells (A549) in 1 h, co-stained for EGFR (green) and cell nuclei (blue) by immunofluorescence, measured by confocal microscopy. b) Increased MSN_tEGFR_ uptake after 1 h in A549 cells normalized to MSN_AVI_ as measured by flow cytometry analysis. ** p = 0.0079; Values given are an average of four independent experiments ± standard error of the mean. Mann-Whitney test. c) Different modes of nanoparticle uptake (*i.e.,* receptor-mediated, blue arrow at (1) and unspecific endocytosis, red arrow at (2)) and endosomal escape, orange arrow at (3) observed in TEM micrographs of A549 cells exposed to MSN_tEGFR_ for 3 h. Cell membrane is shown with the black arrow. Scale bars = 2 μm (upper left), 500 nm for insets 1-3.

### 2.3. Intravenously administered MSNs are deposited in liver and spleen in flank tumor models

Next, we sought to evaluate the targeted uptake of our MSN particles upon systemic delivery in an *in vivo* tumor model. In order to evaluate active targeting efficacy within the same animal, we exploited a double flank tumor model in an immunologically competent mouse strain (C57BL/6) where we subcutaneously inoculated syngeneic clones derived from Lewis lung carcinoma (LLC) cells (**Figure 4A**) or murine melanoma (B16F10) cells that had been genetically engineered to have high or low EGFR expression (*i.e.,* LLC-EGFR^high^ and LLC-EGFR^low^ or B16F10-EGFR^high^ and B16F10-EGFR^low^, respectively).^[22]^ In both settings, cells formed tumors of similar size within two weeks with similar morphology. Fluorescently labeled (ATTO 633) MSNs (with and without targeting ligands) were then systemically applied via retro-orbital intravenous injection and biodistribution of the particles was compared by *in vivo* fluorescence imaging at several time-points up to 48 h post injection. Representative fluorescence images and fluorescence intensity quantifications (Figure 4B&C and Figure S3 and S4A-C) revealed an unspecific increase in biodistribution of all types of particles irrespective of functionalization at the time of injection (t=0) which then slowly decreased back to near baseline-levels over 48 h. To investigate the localization of MSNs on the cellular level, we performed immunofluorescence-based histological analysis of the flank tumors and several internal organs that were previously shown to have enhanced uptake of systemically administered nanoparticles.^[8]^ Both MSN_AVI_ and MSN_tEGFR_ were mainly localized in the liver and spleen with negligible uptake in the flank tumors, lungs, and kidneys (Figure 4D, S4D and S5). Quantification of the immunofluorescence signal per cell nucleus across the different organs confirmed that the localization of the MSNs in the liver was much more pronounced compared to flank tumors and other organs (Figure 4E and Figure S4E). Importantly, we did not observe any difference in the uptake of the particles with regard to whether tumors were derived from EGFR high or low expressing cells. We further confirmed enhanced uptake in the liver compared to LLC flank tumors regardless of MSN type (active *versus* passive) through quantification of nanoparticle-based fluorescence in tissue homogenates of flank tumors and livers (Figure 4F). Together, these data indicate that systemic delivery of actively targeted MSNs does not result in improved uptake into tumors as compared to passively targeted MSNs. Importantly, we found that our nanoparticles did not preferentially localize to the tumors but instead were taken up mostly by the mononuclear phagocyte system of the liver and spleen.

**Figure 4.**
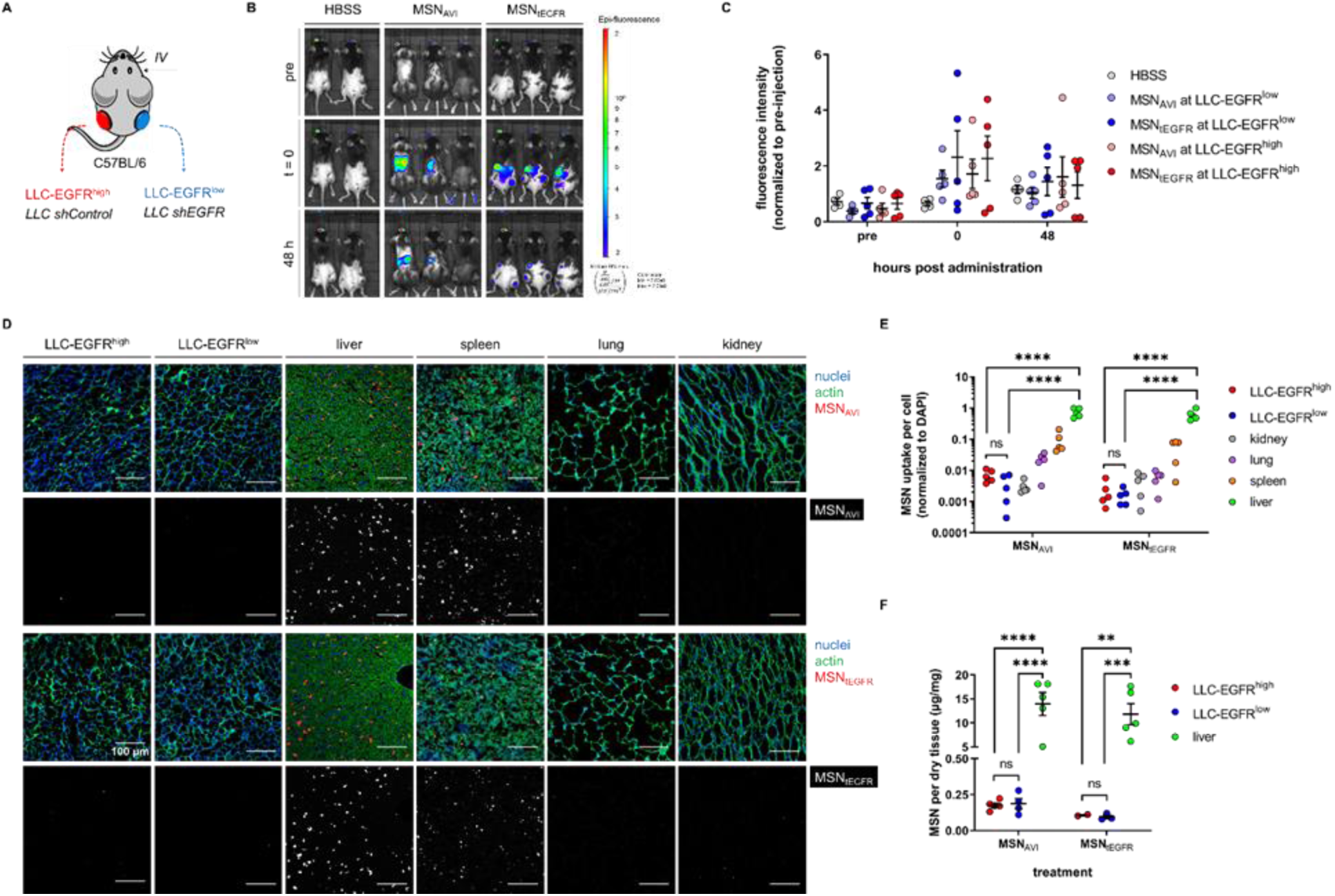
Intravenously administered MSNs are deposited to liver and spleen in a syngeneic lung cancer flank tumor model *in vivo*. a) Schematic representation of the syngeneic double flank tumor-bearing mouse model. LLC clones with different EGFR expression (LLC-EGFR^high^ with high basal EGFR expression and LLC-EGFR^low^ following shEGFR modification; confirmed by Western blot) were injected subcutaneously and developed over two weeks. b) Representative fluorescence images of mice receiving 1 mg of MSN_AVI_ or MSN_tEGFR_ before, immediately after, and 48 h after retro-orbital administration. c) Quantification of the fluorescence intensity obtained from the individual flank tumors of the mice treated with the MSNs over time. Values given are an average of signal obtained from five independent mice at each time point ± standard error of the mean. d) Histological analysis of the biodistribution of intravenously administered MSN_AVI_ and MSN_tEGFR_ in LLC-EGFR^high^ and LLC-EGFR^low^ tumors, livers, spleens, lungs, and kidneys of the mice by confocal microscopy. Nuclear staining (DAPI) is shown in blue, cellular morphology via actin staining (phalloidin) in green and ATTO 633-labeled MSNs in red in the merged image, and in gray in the single channel. Images shown are representative for three different regions from each mice (n = 5 mice treated). Scale bar = 100 μm. e) Quantification of the MSN_AVI_ and MSN_tEGFR_ uptake per nuclei observed in histological analyses in LLC-EGFR^high^ and LLC-EGFR^low^ tumors, kidneys, lungs, spleens, and livers, respectively. **** p < 0.0001, two-way ANOVA, Tukey’s multiple comparisons test, n = 5. f) Quantitative analysis of MSN_AVI_ and MSN_tEGFR_ biodistribution in tissue homogenates of treated animals shows increased uptake in the liver versus either LLC-EGFR^high^ or LLC-EGFR^low^ tumors. Values given are an average of five different samples per MSN type ± SEM. ** p = 0.0022, *** p = 0.0006, **** p < 0.0001, Two-way ANOVA, Sidak’s multiple comparisons test.

### 2.4. Alveolar macrophages engulf intratracheally administered MSNs in a mouse model of lung cancer

One strategy to increase the efficiency of nanoparticle delivery to tumors is through local delivery mechanisms.^[23]^ The lung is considered to be a particularly well-suited organ for local drug delivery as nanoparticles can be delivered via the trachea to the respiratory epithelium of the lung where they are efficiently taken up due to its large surface area, thin epithelium layer, and rich blood supply.^[24]^ Therefore, we next evaluated intratracheal delivery of passively and actively targeted MSNs into the lungs using the previously reported *Kras^LA2^* mutant mouse model for lung cancer.^[25]^ This model displays clinically relevant cancer development compared to tumor cell injection models as tumors develop spontaneously and are heterogeneously distributed (**Figure 5A**). We first evaluated EGFR (Figure 5B) and CCR2 (Figure S6A) overexpression in *Kras^LA2^* mutant lung tumors to confirm that this model is suitable for investigating EGFR- and CCR2-specific targeting via MSN_tEGFR_ (Figure 5) and MSN_tCCR2_ (Figure S6). MSN_AVI,_ MSN_tEGFR_, or MSN_tCCR2_ were intratracheally instilled directly into the lungs of tumor-bearing *Kras^LA2^* mutant mice and the biodistribution of the MSNs was evaluated three days after administration. All MSN types were found to be retained in the lungs of the *Kras^LA2^* mutant mice with no translocation of MSNs to secondary organs (Figure S7). However, particles localized mainly to smaller hyperplastic lesions of tumorous lungs and in the periphery of solid tumors, but no uptake was observed in solid tumors (Figure 5C and Figure S6B). Importantly, high magnification microscopic evaluation did not reveal any obvious differences in cellular uptake of MSN_AVI_, MSN_tEGFR_, and MSN_tCCR2_ on the cellular level (Figure 5D and Figure S6C). The majority of particles were engulfed by alveolar macrophages in both normal and tumorous regions of lungs irrespective of nanoparticle functionalization (Figure 5E and Figure S6D). These data indicate impaired targeting performance of both, actively and passively targeting MSNs to lung tumors when administered via the respiratory surface of the lung due to effective alveolar macrophage clearance of these particles.

**Figure 5.**
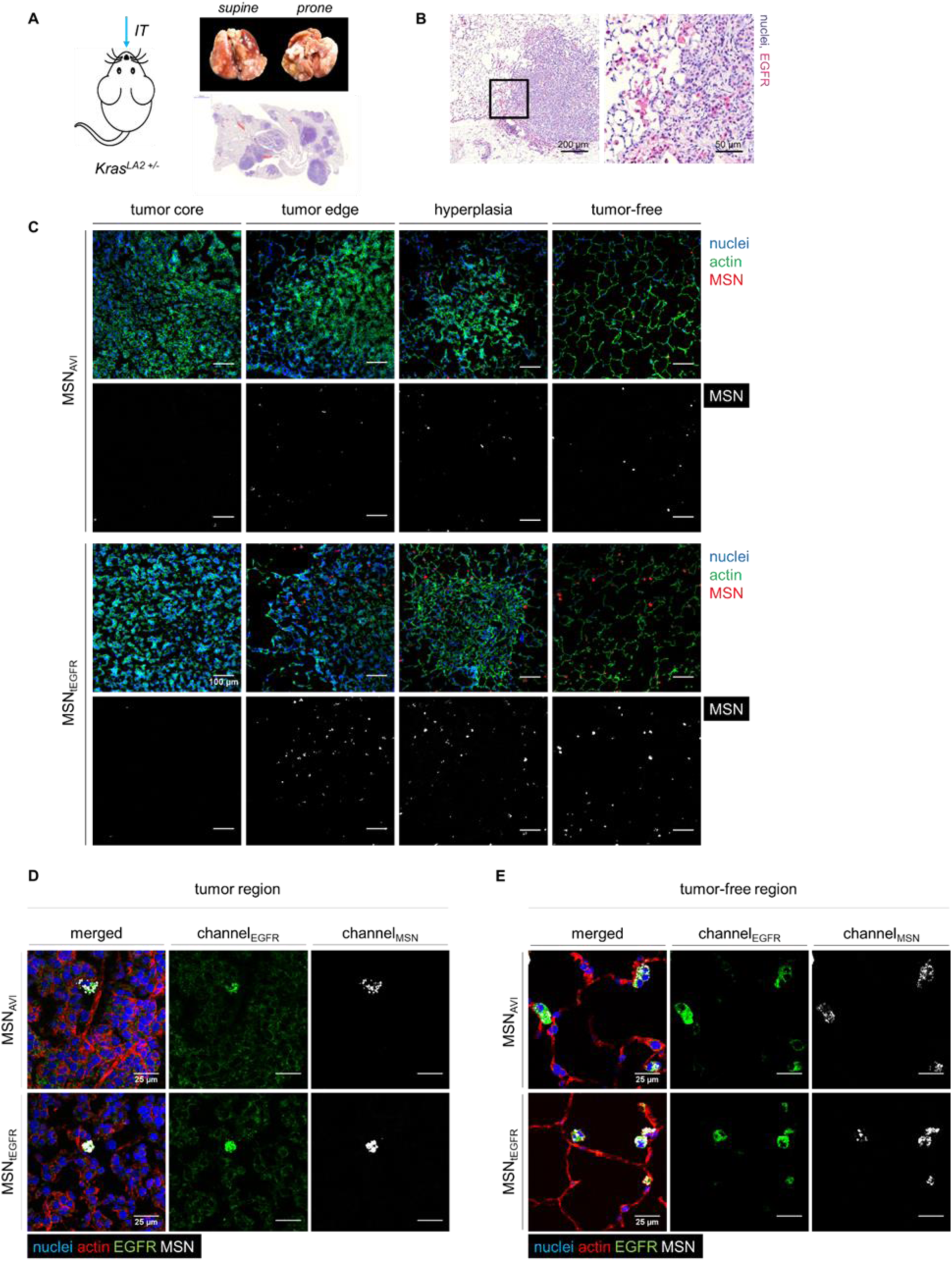
Alveolar macrophages preferentially engulf intratracheally administered MSNs in a *Kras^LA2^* mutant mouse model of lung cancer *in vivo*. a) Representative photographs of a tumorous lung from a *Kras^LA2^* mutant mice in supine and prone positions. b) Immunohistochemistry staining confirming EGFR (pink) overexpression in tumorous regions of *Kras^LA2^* lungs. c) Histological analysis of intratracheally instilled ATTO 633-labeled MSN_AVI_ and MSN_tEGFR_ showing uptake in solid tumor cores *versus* tumor edges, and in hyperplastic or in tumor-free regions of tumorous mouse lungs after 3 days. Nuclear staining (DAPI) is shown in blue, actin staining (phalloidin) in green, and ATTO 633-labeled MSNs in red in the merged images, and in gray in the single channels. Images shown are representative for three different regions from each group of mice (n = 5 per MSN type). Immunofluorescence co-staining for EGFR in d) tumorous *versus* e) tumor-free regions in the mutant lungs treated with ATTO 633-labeled MSN_AVI_ *versus* MSN_tEGFR_. Nuclear staining (DAPI) is shown in blue, cell morphology via actin staining (phalloidin) in red, EGFR staining in green, and ATTO 633-labeled MSNs in gray. Images shown are representative for three different regions from each group of mice (n = 5 per MSN type).

### 2.5. Organ-restricted vascular delivery of MSNs enables specific deposition of nanoparticles in tumors

In order to circumvent clearance by alveolar macrophages, we sought to use an alternative vascular delivery mode which also increases the cross circulatory time of the nanoparticles within the tumor vasculature while minimizing the competing effects such as MPS clearance. Isolated lung perfusion is a clinically established surgical approach which was developed for delivery of higher doses of chemotherapy to the lung through isolation of the pulmonary circulation from systemic circulation.^[15]^ While the procedure itself is safe, it has not yet seen widespread clinical application due to damage to the neighboring healthy tissue from chemotherapeutics. Therefore, we sought to leverage our pH-responsive nanoparticles by delivering them using a modified isolated lung perfusion procedure. The combination of isolated lung perfusion with environmentally-responsive or stimuli-responsive nanoparticles could permit controllable delivery of chemotherapeutics selectively to tumorous tissue.

Owing to the difficulties in performing such a surgery in a small animal model of lung cancer, we tested the feasibility of this approach in an *ex vivo* model to simulate the surgical conditions of isolated lung perfusion. We explanted tumorous lungs from healthy or *Kras^LA2^* mutant mice and introduced actively or passively targeting MSNs for 3 h at controlled flow rates using an *ex vivo* system containing ventilation and perfusion which we have previously described (**Figure 6A**).^[26]^ Our first data using organ-restricted vascular delivery (ORVD) indicated that nanoparticles were homogeneously distributed in healthy mouse lungs and located with close proximity to endothelial cells (CD31 positive), suggesting their retention inside healthy blood vessels (Figure S8A). On the other hand, we observed enhanced delivery of MSN_AVI_, MSN_tEGFR_, and MSN_tCCR2_ into the cores of solid lung tumors in a reproducible manner, regardless of their surface functionalization for active targeting (Figure 6B). Similar to other *ex vivo* murine lung setups used for pharmacologic analysis, we did not observe any significant cell death (*i.e.,* cleaved caspase-3) over the 3 h exposure time ^[27]^, indicating that neither the nanoparticles nor surgical ORVD technique induced cytotoxic side-effects. Additionally, we did not observe uptake of nanoparticles by lung-resident macrophages (Figure S8B) but did observe efficient cellular uptake of the nanoparticles by cancer cells located in solid tumor cores as demonstrated by TEM analysis (Figure 6C). In lungs exposed to MSN_tEGFR_ via ORVD, nanoparticles localized to tumor cells with lamellar bodies, validating their distal epithelial origin (Figure 6C). Taken together, these results demonstrate that our organ-restricted approach results in enhanced cellular uptake of actively and passively targeting nanoparticles in solid lung tumors. ORVD may thus represent a useful strategy to overcome some of the existing physicochemical and biological barriers limiting nanoparticle delivery efficacy to solid tumors of the lung.

**Figure 6.**
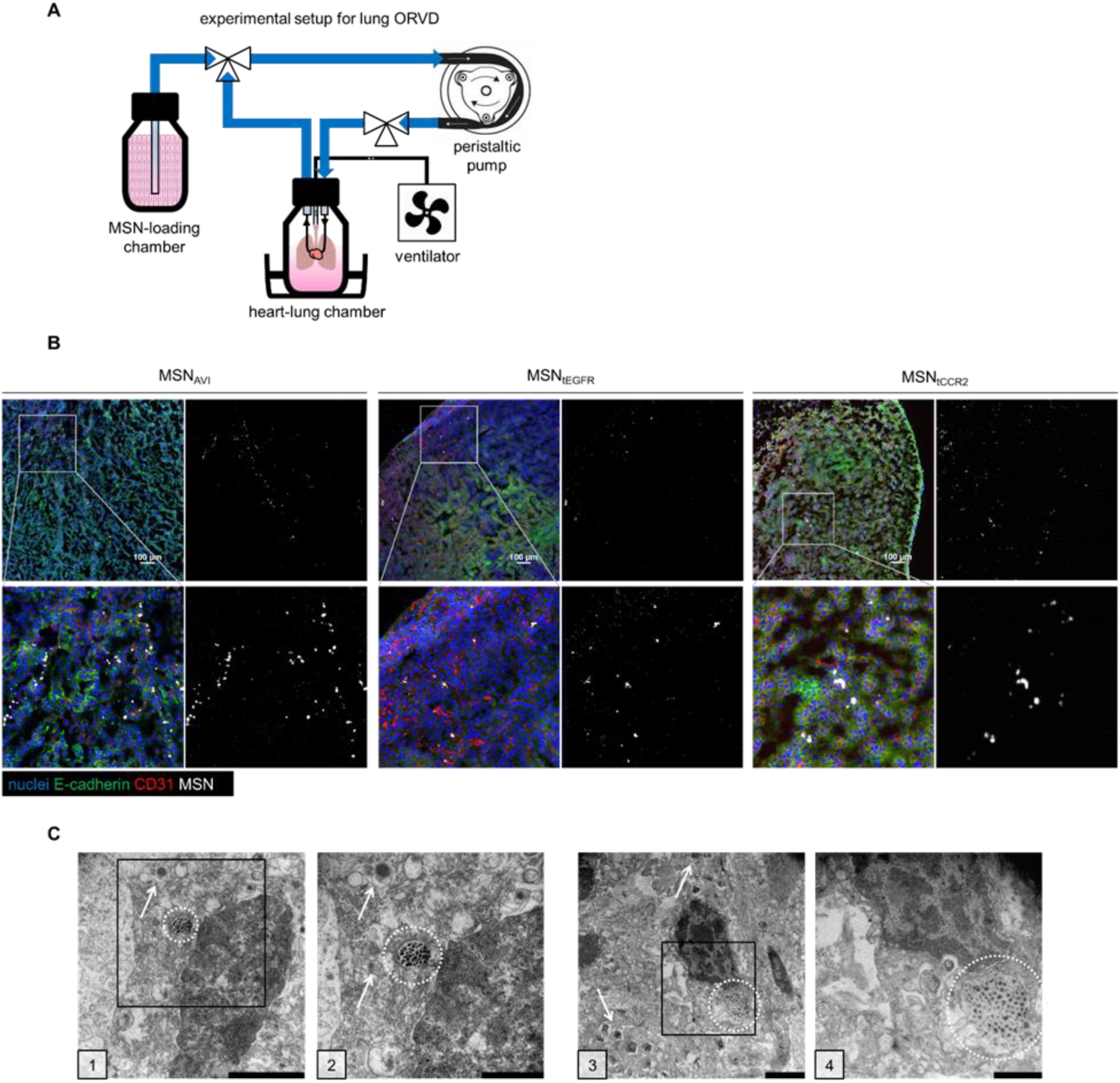
Organ-restricted vascular delivery of MSNs results in enhanced cellular uptake of the particles in solid lung tumors *ex vivo*. a) Schematic representation of the organ-restricted vascular delivery (ORVD) setup with *ex vivo* lung perfusion and ventilation. Murine heart and lung blocs were mounted in a custom-designed bioreactor with temperature control (37°C) and connected to an external chamber containing MSNs loaded perfusate which were instilled at hypoperfusion rates (0.5 mL/min). Lungs were ventilated at 100 strokes/min with a 100 μL stroke volume. b) Representative histological analysis showing uptake of MSNs into solid tumors from the *Kras^LA2^* lungs following ORVD; MSN_AVI_, MSN_tEGFR_, and MSN_tCCR2_ (gray in each panel), nuclei stained with DAPI (blue), epithelial cells labeled with E-Cadherin (green) and endothelial cells labeled with CD31 (red) shown in merged and corresponding single channel images. Scale bar in the top row = 100 μm. c) Intracellular MSN_tEGFR_ (highlighted in white dashed circles) in *Kras^LA2^* mutant solid lung tumor cores after 3 h of ORVD, visualized with TEM. Lamellar bodies (white arrows) indicate uptake in cells of alveolar type II origin; scale bars are 2 μm for image panels 1 and 3, 1 μm for image panels 2 and 4.

## 3. Conclusion

Despite rapid and exciting progress in the design and synthesis of increasingly sophisticated nanomedicines, overcoming the physical and biological barriers of specific organs has remained a primary challenge upon administration *in vivo*. Additionally, recent evidence suggests that previous reports may have overestimated the contribution of the EPR to directing nanoparticles to tumors and that additional active cellular processes, such as transcytosis, also promote extravasation of nanoparticles into tumors.^[16]^ Many of these experiments have demonstrated proof-of-concept nanoparticle delivery in immunocompromised animals thereby underestimating deposition to immunological active off-target organs upon systemic administration, which remains a major challenge.

Therefore, in order to circumvent systemic effects associated with low efficacy of nanoparticle delivery, we developed a novel organ-restricted delivery strategy to directly administer nanoparticles into the target organ vasculature which allows recirculation. We also show that the ORVD technique promotes their extravasation from the deranged tumor vasculature into the core of solid tumors possibly via the recently demonstrated transcytosis pathway.^[16]^ Encouragingly, we observed specific cellular uptake of nanoparticles within the core of solid lung tumors while nanoparticles were retained within the capillaries in regions with healthy tissue. This observation is fully in line with the recent work showing preferential activation of transcytotic transport of particles across the tumor vasculature. We did not observe any noticeable differences in cellular uptake between active or passively targeting nanoparticles using ORVD suggesting that the delivery method is the primary hurdle to obtain effective delivery to the tumor site.

Lung cancer remains one of the most challenging cancers to treat and its prevalence is increasing globally.^[28]^ In addition to the fact that it is often diagnosed at advanced stages when tumors are beyond surgical treatment due to their larger size or their location, the lung is also a major site of metastasis and metastasized cancers are some of the most challenging cancers to treat.^[29]^ Even at early detection stages, some small tumors remain inoperable and cannot be removed due to their location centrally or due to reduced pre-operative lung capacity. Additionally, patients with reduced lung capacity might not be able to undergo wedge or lobe resection. Overall survival for lung cancer is around 15% after 5 years. Currently, only 20% of lung cancer patients are eligible for resections to remove large tumors due to the location of the tumor and potential co-morbidities which may present complicating factors ^[30]^. Resection is often followed by radiation and chemotherapeutics to address hyperplastic regions and to avoid recurrence after surgery.^[31]^ The majority of lung cancer chemotherapeutics are administered systemically and are associated with negative side effects (*e.g.,* anemia), and are particularly toxic to the kidney, nervous system, and heart.^[32]^ A substantial number of patients have to end or pause the treatment prematurely due to negative side effects. Recently, nanoparticles have emerged as a potential option to circumvent some of the problems associated with systemic administration of chemotherapeutics, but numerous challenges remain and therefore the efficacy of nanoparticle-mediated drug delivery to solid tumors has been limited.^[33]^

In this study, we designed and compared cell-specific targeting efficacies of actively and passively targeted pH-responsive mesoporous silica nanoparticles for treatment of lung cancer. The mesoporous silica nanoparticles we utilize here have several benefits which make them an attractive nanoparticle platform for treating cancer: 1) they are inert and can be loaded with virtually any cargo smaller than their pore size (*i.e.,* 4 nm), making it compatible with a range of existing and emerging chemotherapeutics and other compounds, 2) contain a pH cleavable linker, restricting delivery of their cargo to cells within the acidic tumor microenvironment or after cellular uptake in the endo-lysosome, 3) can readily be functionalized for active targeting, and 4) are cytocompatible.^[17, 34]^ Overall, MSNs are highly adaptable and can be fine-tuned for each particular cancer microenvironment.

For active targeting studies, we explored a combinatorial approach to address complementary compartments of solid tumors:^[35]^ the parenchyma via EGFR targeting and the stroma via CCR2 targeting, since these two surface receptors have been previously shown to be overexpressed in lung cancer and associated with poor prognosis of the disease.^[18–19, 36]^ As has been shown in previous studies for MSNs and other nanoparticles ^[2, 37]^, MSNs which contained active targeting ligands were taken up by cells at a significantly higher rate than passively targeted nanoparticles *in vitro*. However, we did not observe any preferential uptake of actively targeted nanoparticles when they were administered intravenously *in vivo*.

In order to minimize the likelihood that our active targeting strategy failed due to differential uptake between tumor types, we used two different *in vivo* flank tumor models in this study: 1) a murine lung carcinoma line (LLCs) to mimic primary lung tumors and 2) melanoma line (B16F10) to mimic melanoma metastasis in the lung. Furthermore, we included an internal control for each animal by generating paired flank tumors on the left and right side of each animal using two different clones which were genetically engineered to differentially express high or low amounts of EGFR, respectively. Of note, we did not observe any differences in tumor uptake for active or passively targeting MSNs, nor between tumors derived from tumorous cells engineered for high or low levels of EGFR. Finally, our flank models were syngeneic and thus generated in immunocompetent mouse models, making them relevant to the clinical scenario where a number of immune associated biological barriers play a role in limiting nanoparticle efficacy.^[8, 33]^ In all of these flank models, we found that the majority of nanoparticles were taken up in the liver and spleen, independent of the presence or absence of a targeting ligand, which is in line with some previous work.^[7a]^

Local administration of chemotherapeutics and nanoparticles via the airways has been previously attempted for lung cancer.^[2, 13]^ Therefore, we next tested our nanoparticles via intratracheal administration in the *Kras^LA2^* mutant mouse model, which spontaneously forms lung tumors over a period of several months.^[25]^ This model is highly representative of the clinical scenario as the tumors are heterogeneous in size and distributed throughout the lung. Furthermore, *Kras* is the most frequent oncogene driver mutation in lung cancer ^[38]^ for which no approved drugs exist. After lung tumors were allowed to develop in these animals, we administered the nanoparticles intratracheally. Although, we did not observe any nanoparticle localization in other internal organs 3 days post-instillation, the overwhelming majority of MSNs were taken up nonspecifically *in vivo* by alveolar macrophages in both normal and tumorous lungs. There was no obvious difference in uptake between active and passively targeting nanoparticles. Intriguingly, our *in vitro* experiments showed higher uptake of the receptor-targeted nanoparticles, which suggests that the loss of targeting *in vivo* is a result of several additional factors such as the inherent capacity of the MPS to efficiently clear systemically or locally delivered particles. Our data are partly in contrast to other studies that observed enhanced tumor delivery using active *versus* passively targeted nanoparticles *in vivo.*^[39]^ Differences between our findings and those of other groups may be due to the targeting ligands chosen, nanoparticle shape and composition, the animal model used, target organ/tumor architecture, and how tumor uptake and drug delivery efficacy was assessed.^[8]^

Our work demonstrates the potential of administering nanoparticles in an organ-restricted manner to increase delivery and cellular uptake into solid lung tumors. The ORVD approach represents a novel strategy for locally delivering nanoparticles into the lung or other highly vascularized organs. By using controlled perfusion and recirculation of nanoparticles we promote extravasation from leaky, tumor-associated blood vessels and subsequently promote retention and cellular uptake in targeted tumor cells. As we did not observe any significant cellular damage in our ORVD setup, this indicates that the cellular uptake we observed was likely due to either the EPR effect or increased transcytosis in tumorous regions and not due to the induction of vascular damage in our *ex vivo* setup. We chose to perform our initial ORVD experiments in an *ex vivo* model due to the difficulties in performing isolated lung perfusion in murine animals *in vivo*. *Ex vivo* models and evaluation of potential therapies are an emerging research area and provide a unique opportunity to perform preclinical testing for situations which are difficult to first mimic *in vivo*. ^[27, 40]^ While these data provide proof-of-principle evidence that ORVD is applicable for nanoparticle-based targeting of solid lung tumors, it will be important in future studies to validate this approach in larger animals using chemotherapeutic-loaded nanoparticles *in vivo* ultimately aiming at the clinical application of this concept in cancer patients.

Isolated lung perfusion uses extracorporeal circulation techniques to isolate the pulmonary vasculature from the systemic circulation. It has already been used for delivering lung cancer chemotherapeutics with the intent of increasing local delivery concentrations and reducing systemic toxicity.^[14]^ However, despite the fact that the surgical approach is safe, one of the major limiting factors for more widespread use of this surgical technique has been the toxicity associated with the surrounding healthy tissue which is also exposed to chemotherapeutics^[14]^. Smart, stimuli-responsive nanoparticles, such as those used in this study, offer the unique advantage of controlled and selective release of drugs into the tumor microenvironment. Furthermore, application of nanoparticles under controlled flow parameters during ORVD may allow for further fine-tuning of delivery to optimize nanoparticle uptake under EPR conditions. In addition, it will be of interest to further explore other nanoparticle formulations, including those which showed safety but no efficacy in clinical trials when administered systemically.

In summary, we have shown that the extended recirculation of actively and passively targeted mesoporous silica nanoparticles in an organ-restricted fashion via isolated pulmonary perfusion results in enhanced localization of the nanoparticles in solid lung tumors. These results bring optimism to be able to offer a new treatment option for patients with inoperable lung tumors and those who are unable to cope with chemotherapy due to negative side effects. Our findings represent a clinically relevant and promising strategy to direct the nanoparticles to solid tumors in highly vascularized tissues for an enhanced therapeutic efficacy.

## 4. Experimental Section

### Synthesis of core-shell functionalized mesoporous silica nanoparticles (MSN-SH_in_-NH_2,out_)

A mixture of tetraethyl orthosilicate (TEOS, 1.63 g, 7.82 mmol), mercaptopropyl triethoxysilane (MPTES, 112 mg, 0.48 mmol) and triethanolamine (TEA, 14.3 g, 95.6 mmol) was heated under static conditions at 90°C for 20 min in a polypropylene reactor. Then, a solution of cetyltrimethylammonium chloride (CTAC, 2.41 mL, 1.83 mmol, 25 wt% in H_2_O) and ammonium fluoride (NH_4_F, 100 mg, 2.70 mmol) in H_2_O (21.7 g, 1.21 mmol) was preheated to 60°C, and rapidly added to the TEOS solution. The reaction mixture was stirred vigorously (700 rpm) for 20 min while cooling down to room temperature. Subsequently, TEOS (138.2 mg, 0.922 mmol) was added in four equal increments every three min. After another 30 min of stirring at room temperature, TEOS (19.3 mg, 92.5 µmol) and aminopropyl triethoxysilane (APTES, 20.5 mg, 92.5 µmol) were added to the reaction. The resulting mixture was then allowed to stir at room temperature overnight. After addition of ethanol (100 mL), the MSNs were collected by centrifugation (7000 rcf, for 20 min) and re-dispersed in absolute ethanol. The template extraction was performed by heating the MSN suspension under reflux (90°C, oil bath temperature) for 45 min in an ethanol solution (100 mL) containing ammonium nitrate (NH_4_NO_3_, 2 g), followed by 45 min heating under reflux in a mixture of concentrated hydrochloric acid (HCl, 10 mL) and absolute ethanol (90 mL). The mesoporous silica nanoparticles were collected by centrifugation and washed with absolute ethanol after each extraction step.

### Synthesis of MSN_COOH_

A large excess of oxalic acid (10 mg, 110 µmol) was dissolved in 2 mL water and activated with EDC (18 µL, 102 µmol) and a catalytic amount of sulfoNHS (1 mg) for 10 min at room temperature. The premixed solution was added dropwise to 100 mg MSN-SH_in_- NH_2,out_ particles dissolved in 15 mL ethanol under vigorous stirring. The mixture was stirred at room temperature overnight. Afterwards the solution was centrifuged at 7000 rcf for 10 min, washed two times with ethanol and redispersed in 10 mL ethanol.

### Synthesis of MSN_AK_

25 mg of MSN_COOH_ were diluted in 15 mL ethanol. Subsequently, 10 µL N- (3-dimethylaminopropyl)-N’-ethylcarbodiimide hydrochloride (EDC, 57 µmol) and 3.1 mg of N-hydroxysulfosuccinimide (sulfo-NHS, 14.3 µmol) were added and the mixture was stirred for 15 min at room temperature. A premixed solution containing of 3.5 mg 3,9-bis(3-aminopropyl)- 2,4,8,10-tetraoxaspiro-[5,5′]-undecane AK-Linker (13 µmol) in 3 mL of a 1/1 mixture ethanol/DMSO were added dropwise over a period of 10 min and the resulting solution was stirred over night at room temperature. The functionalized MSN_AK_ particles were separated by centrifugation (7000 rcf, 20 min), washed two times with ethanol and redispersed in 15 mL ethanol.

### Synthesis of MSN_Biotin_

A premixed solution of 1 mg biotin (4.1 µmol), 1 µL N-(3-dimethylaminopropyl)-N’-ethylcarbodiimide hydrochloride (EDC, 5.7 µmol) and 1.2 mg N-hydroxysulfosuccinimide (sulfo-NHS, 5.7 µmol) were added to 10 mg of MSN_AK_ particles in 5 mL ethanol and stirred overnight at room temperature. After centrifugation (7000 rcf, 20 min) and washing two times with ethanol, MSN_Biotin_ particles were separated by centrifugation and redispersed in 5 mL ethanol.

### Dye-labeling of MSN_Biotin_

1 mg MSN_Biotin_ were diluted in 1 mL ethanol, and 1 µL ATTO 633- or ATTO 488-maleimide (0.5 mg/mL in DMF) was added. The mixture was reacted for 12 h overnight in the dark. Afterwards the particles were centrifuged (7000 rcf, 5 min), washed twice with ethanol and resuspended in HBSS buffer to give a 1 mg/mL particle concentration.

### Synthesis of MSN_AVI_

After centrifugation (14000 rpm, 4 min) the loaded or non-loaded residue (MSN_Biotin_) was redispersed in a solution containing of 1 mg avidin from egg white in 1 mL HBSS buffer solution and stored for 1 h at room temperature in the dark without stirring. The resulting suspension was then centrifuged (5000 rcf, 4 min, cooled) and washed several times with buffer solution. Subsequently, the particles were finally redispersed in 1 mL of the corresponding buffer solution and used for the following experiments.

### Addition of the targeting ligands to synthesize MSN_tEGFR_ and MSN_tCCR2_

1 mg of cargo-loaded and/or dye-labeled MSN_AVI_ particles were centrifuged (5000 rcf, 4 min, cooled) and redispersed in 500 µL HBSS buffer solution. In the meantime, 50 µL of the corresponding targeting ligand (GE11 or ECL1i) dissolved in bi-distilled water (100 µg/mL) were added to 200 µL HBSS and 0.2 mg 2-iminothiolan hydrochloride (1.5 µmol). The mixture was reacted for 1 h at room temperature without stirring. Subsequently, 0.3 mg of the hetero-bifunctional PEG-linker mal-PEG3000-NHS was added and the mixture was allowed to react for 1 h at room temperature. The activated PEG-targeting ligand was then added to the MSN_AVI_ particle solution, activated for 1 h at room temperature without stirring. Subsequently, 0.3 mg of the hetero-bifunctional PEG-linker mal-PEG3000-NHS was added and the mixture was allowed to react for 1 h at room temperature. The activated PEG-targeting ligand was then added to the MSN_AVI_ particle solution, reacted for 1 h, centrifuged (5000 rcf, 4 min, cooled) and washed three times with HBSS. 1 mg of MSN_tEGFR_ and MSN_tCCR2_ were redispersed in 1 mL HBSS, respectively.

### Characterization methods

Dynamic light scattering (DLS) and zeta potential measurements were performed on a Malvern Zetasizer-Nano instrument equipped with a 4 mW He-Ne laser (633 nm) and an avalanche photodiode detector. DLS measurements were directly recorded in diluted colloidal aqueous suspensions of the MSNs at a constant concentration of 0.5 mg/mL for all sample solutions. Zeta potential measurements were performed using the add-on Zetasizer titration system (MPT-2), based on diluted NaOH and HCl as titrants. For this purpose, 0.5 mg of the MSN sample was diluted in 10 mL bi-distilled water. Transmission electron microscopy (TEM) was performed at 300 kV on an FEI Titan 80-300 equipped with a field emission gun. For sample preparation, the colloidal solution of MSNs was diluted in absolute ethanol, and one drop of the suspension was then deposited on a copper grid sample holder. The solvent was allowed to evaporate. Thermogravimetric analyses (TGA) of the extracted bulk samples (approximately 10 mg) were recorded on a Netzsch STA 440 C TG/DSC. The measurements proceeded at a heating rate of 10 °C/min up to 900°C, in a stream of synthetic air of about 25 mL/min. Nitrogen sorption measurements were performed on a Quantachrome Instrument NOVA 4000e at −196°C. Sample outgassing was performed for 12 h at a vacuum of 10 mTorr at 120°C or room temperature. Pore size and pore volume were calculated with an NLDFT equilibrium model of N_2_ on silica, based on the desorption branch of the isotherms. In order to remove the contribution of the interparticle textural porosity, pore volumes were calculated only up to a pore size of 8 nm. A BET model was applied in the range of 0.05 – 0.20 p/p_0_ to evaluate the specific surface area. Infrared spectra were recorded on a ThermoScientific Nicolet iN10 IR-microscope in reflection-absorption mode with a liquid-N_2_ cooled MCT-A detector. For time-based release experiments of propidium iodide, the loaded and avidin-capped particles were redispersed in the corresponding buffer solutions (pH = 7 and pH = 5) and stored at 37°C on a thermo shaker. After certain time-points (4 h, 24 h, 48 h) the particles were centrifuged (5000 rcf, 4 min, cooled) and the supernatant was measured on a UV/VIS Thermo Scientific NanoDrop 2000c system. For scanning electron microscopy, 5 µl of 1 mg/mL MSN_tEGFR_ in 100% ethanol solution were pipetted onto a 12.5 mm aluminum stub and left to air dry. Samples were subsequently sputter coated with 2 nm Pt/Pd (80/20) in a Quorum Q150T ES turbo pumped sputter coater and examined with the secondary electron detector at 1.5 kV in a Jeol JSM-7800F FEG-SEM.

### Human tissue

The staining with human tissue was approved by the Ethics Committee of the Ludwig-Maximilians-University Munich, Germany (LMU, project no. 455-12). All samples were provided by the Asklepios Biobank for Lung Diseases, Gauting, Germany (project no. 333-10). Written informed consent was obtained from all subjects.

### Cell culture

The human non-small-cell lung cancer cell lines, A549 and H520, and the mouse alveolar macrophage cell line, MH-S, were obtained from the ATCC (American Type Culture Collection). A549 and H520 cells were maintained in DMEM medium supplemented with 10% FBS and 1% Pen/Strep. MH-S cells were maintained in RPMI 1640 medium supplemented with 10% FBS and 1% Pen/Strep. MH-S cells were further supplemented with 1 mM sodium pyruvate, 10 mM HEPES, and 50 µM β-mercaptoethanol (all AppliChem). All cells were grown at 37°C in a sterile humidified atmosphere containing 5% CO_2_.

### Immunocytofluorescence

A549 and MH-S cells which were grown on coverslips were exposed to ATTO 633-labeled MSNs for 1 h. Afterwards, the cells were washed three times with PBS, then once with NaCl (0.15 M, pH 3.0), and then three times with PBS again. Cells were fixed with 70% ethanol and permeabilized with 0.1% Triton-X. After another PBS wash, cells were incubated with Roti-Block for 1 h at room temperature. Then, A549 cells were stained with EGFR antibody (Abcam, ab52894) whereas MH-S cells were stained with CCR2 antibody (Novus Biologicals, NB110-55674) overnight at 4°C. The following day, the cells were incubated with the Alexa Fluor secondary antibodies for 1 h at room temperature, washed with PBS, incubated with DAPI for 10 min for nuclear staining, and then mounted with fluorescent mounting medium (Dako).

### Western blotting

A549, H520, and MH-S cells were lysed in RIPA buffer (50 mM Tris-HCl, pH 7.5, 150 mM NaCl, 1% NP-40, 0.5% sodium deoxycholate, 0.1% SDS) supplemented with cOmplete protease inhibitor cocktail. Protein content was determined using the Pierce BCA protein assay kit (Thermo Scientific). For Western blot analysis, equal amounts of protein were subjected to electrophoresis on 10% SDS-PAGE gels and blotted onto PVDF membranes (Bio-Rad). Membranes were treated with antibodies using standard Western blot techniques. The ECL Plus detection reagent (GE Healthcare) was used for chemiluminescent detection and the membranes were analyzed with the ChemiDoc XRS+ (Bio-Rad).

### Flow cytometry

5×10^5^ A549 or MH-S cells were plated on 6 well plates and incubated overnight. The next day, the cells were exposed to ATTO 488- or ATTO 633-labeled MSNs for 1 h. Afterwards, the cells were washed three times with PBS, once with NaCl (0.15 M, pH 3.0), and then three times with PBS again to create a final cell suspension. Samples were then analyzed by flow cytometry (BD LSRII). MSN uptake in different cell types was quantified by the median fluorescence signal collected in the Alexa Fluor 488 or 647 channels.

### Transmission electron microscopy

Samples were fixed in 2% paraformaldehyde and 2% glutaraldehyde in 0.1 M Sorensen phosphate buffer. After fixation, the specimens were rinsed in buffer, post-fixed in 1% osmium tetroxide, dehydrated using graded acetone solutions and, embedded in Polybed 812 epoxy resin. Ultrathin sections were cut using Leica EM UC7, mounted on Maxtaform H5 copper grid. The sections were stained with 2% uranyl acetate and 1% lead citrate. The sections were then analyzed in FEI Tecnai BioTwin 120 kV microscope.

### Animal models

#### Syngeneic flank tumor models

C57BL/6 mouse Lewis lung carcinoma (LLC) and B16F10 skin melanoma cells were obtained from the NCI Tumor Repository (Frederick). For RNA interference, the following proprietary lentiviral shRNA pools were obtained from Santa Cruz Biotechnology (Palo Alto): random control shRNA (shC, sc-108080), GFP control (sc-108084), anti-EGFR-shRNA (sc-29302-V), and stable transfections of the LLC and B16F10 cells were generated as described previously.^[22]^ C57BL/6 mice were obtained from Jackson Laboratories (Bar Harbor) and were bred at the Center for Animal Models of Disease of the University of Patras, Greece. Experiments were approved a priori by the Veterinary Administration of the Prefecture of Western Greece, and were conducted according to Directive 2010/63/EU. Experimental mice were sex-, weight-, and age-matched. For induction of solid tumors, mice were anesthetized using isoflurane inhalation and received subcutaneous injections of 100 µL PBS containing 0.5 x 10^6^ LLC or B16F10 clones.

#### Transgenic lung cancer model

129S/Sv-Kras^tm3Tyj^/J (*Kras^LA2^*) mutant mice were obtained from the Jackson Laboratory, USA, and cross-bred with FVB-NCrl WT females obtained from Charles River Laboratories, Germany, for over seven generations. Animals were kept in rooms maintained at constant temperature and humidity with a 12/12 h light/dark cycle and were allowed food and water *ad libitum*. Animal experiments were carried out according to the German law of protection of animal life and were approved by an external review committee for laboratory animal care.

### In vivo biodistribution studies

#### Intravenous application

Two weeks after subcutaneous inoculation of EGFR-high and EGFR-low LLC and B16F10 tumor clones, 1 mg ATTO 633-labeled MSN_AVI_ or MSN_tEGFR_ was applied to each mouse retro-orbitally. The mice were sacrificed with an overdose of isoflurane three days after the administration. *In vivo* live animal imaging experiments were carried out to analyze the pharmacokinetics and *ex vivo* organ distribution of ATTO 633-labeled MSNs. Fluorescence imaging of living mice was done using an IVIS Lumina II imager (Perkin Elmer, Santa Clara, CA). Mice were anesthetized using isoflurane and serially imaged at various time-points: before, immediately at, 3, 6, 24, and 48 h post-injection of MSNs. Retro-orbital venous sinus injection, comparable to tail-vein injection, was used, in order to avoid potential animal distress and/or retention of significant amounts of dose in the tail. The images were acquired and analyzed using Living Image v4.2 software (Perkin Elmer, Santa Clara, CA). Flank tumors were selected as specific regions of interest and photon flux within these regions were measured. *Intratracheal application:* 12 week-old *Kras^LA2^* mutant mice were intratracheally instilled with ATTO 633-labeled MSN_AVI_, MSN_tEGFR_, and MSN_tCCR2_, as previously described ^[34]^. Three days post-instillation, the mice were sacrificed with an overdose of ketamine (188.3 mg/kg) and xylazin hydrochloride (4.1 mg/kg) (bela-pharm). Lung lobes from each group (n=5 mice per group) were excised and prepared for cryoslicing.

#### Organ-restricted vascular delivery

Heart-lung blocks from WT and *Kras^LA2^* mutant mice were extracted and placed into the *ex vivo* perfusion/ventilation system as described by Bölükbas *et al.*^[26]^ and schematically depicted in Fig 5A. The system was protected from light for all of the experiments containing the fluorescent MSNs. The *ex vivo* heart-lung blocks were submerged in cell culture media supplemented with 10% FCS and 1% Pen/Strep in the internal incubation chamber kept at 37°C and were mechanically ventilated with a respiratory frequency of 100 strokes/minute and 100 μL stroke volume throughout the 3 h nanoparticle exposure. The blocks were perfused with PBS for 10-15 min before intra-arterial administration of the particles via the pulmonary trunk. 400 µL of 1 mg/mL concentrated ATTO 633-labeled MSN suspension was dispersed in 20 mL cell culture media in the external chamber protected from light. The MSN solution from the external chamber was fed into the heart-lung block for 10 min at 0.5 mL/min flow rate using a peristaltic pump for the initial loading of the system with 100 µg MSNs. Then, the feeding was stopped and the particle suspension was re-circulated through the inner loop of the system for an additional 160 min. After the exposure, the lungs were filled with OCT for cryosectioning and histological observation.

#### Immunohistochemistry

Lung tumor specimens from human and *Kras^LA2^* mutant mice were placed in 4% (w/v) paraformaldehyde (PFA) overnight at 4°C and processed for paraffin embedding. 3 μm thick paraffin sections were sliced with the microtome (Zeiss Hyrax M 55) and placed on superfrost plus adhesion slides. De-paraffinized sections were subjected to quenching of endogenous peroxidase activity using a mixture of methanol/H_2_O_2_ for 20 min, followed by antigen retrieval in a de-cloaking chamber. From this step on, the slides were washed with tris-buffered saline with Tween-20 (TBST, 20 mM Tris, 0.8% NaCl, 0.02% Tween-20, pH 7.6 adjusted with HCl) after each incubation with the reagents throughout the procedure. The sections were incubated first with Rodent Block M (Zytomed Systems) for 30 min and then with the primary antibody, *i.e.,* EGFR (Cell Signaling, D38B1 for human, Abcam, ab52894 for mouse), CCR2 (Novus Biologicals, NB110-55674), or IgG control for 1 h. The cuts were then incubated with Rabbit on Rodent AP-Polymer for 30 min, which was followed by Vulcan Fast Red AP substrate solution (both Biocare Medical) incubation for 10-15 min. Sections were counterstained with hematoxylin (Carl Roth) and dehydrated respectively in consecutively grading ethanol and xylene (both Appli-Chem) incubations. Dried slides were mounted in Entellan and visualized with a slide scanner.

#### Histological preparations and immunofluorescence imaging

For the intravenous systemic delivery experiment, internal organs as well as flank tumors were dissected and placed in 4% PFA overnight after which the suspension medium was exchanged to PBS. Representative parts of the organs were frozen in OCT and kept at −80°C. For the lungs obtained from the intratracheal delivery as well as the ORVD experiments, the airways were immediately filled with OCT by intratracheal administration. Later, the lobes were separated, transferred into cryomolds, and covered with additional OCT. Samples were left to freeze on dry ice and then stored at −80°C. For both experiments, 5 μm thick cryo-sections were sliced with a cryostat (Zeiss Hyrax C 50) and placed on superfrost plus adhesion slides. Immediately before staining, all cryo-sections were fixed with 4% PFA for 10 min, then washed with PBS, and permeabilized with 0.5% Triton-X. The sections were incubated with Roti-Block for 1 h at room temperature, and then with the primary antibody at 4°C overnight; *i.e.,* EGFR (Abcam, ab52894) and CCR2 (Novus Biologicals, NBP1-48338). Afterwards, the sections were washed with PBS, incubated with Alexa Fluor 488 secondary antibody for 1 h at room temperature. After another PBS wash, the sections were finally stained with DAPI. In case phalloidin staining was used, the sections were first incubated with phalloidin for 45 min and then with DAPI for 10 min at room temperature directly after the fixation and washing step. The sections were mounted using fluorescence mounting medium (DAKO) and analyzed using confocal microscopy (LSM710, Carl Zeiss). Quantification of the cellular uptake of the MSNs in the tissues was conducted using the IMARISx64 software (version 7.6.4, Bitplane).

#### Fluorescence dosimetry of MSNs in organ homogenates

The dose of ATTO 633-labeled MSNs in the flank tumors and livers was determined with quantitative fluorescence analysis similar to the validated method described by Rijt *et al.*^[34]^ Briefly, aliquots of the tissue were dried at low power setting in a microwave oven (SEVERIN, MW7803; 30% power; 270 Watt) until no change of tissue mass was observed anymore. Aliquots of dried tumor and liver tissue (ca. 10 mg) were diluted by 1:90 (w/v) and 1:60 with PBS, respectively (*i.e.,* 1 mg of dried tissue was diluted by 89 and 59 µL PBS, respectively). The diluted samples were mechanically homogenized with a high-performance disperser (T10 basic ULTRA-TURRAX^®^) on ice until no tissue pieces were visible anymore (ca. 3-5 min with short breaks to avoid undue heating of the samples). Residual tissue was rinsed off the disperser using 200 μL of PBS. Samples were vortexed immediately prior to pipetting four 75 μL aliquots (quadruple determination) from each of the samples into a black 96-well plate for quantitative fluorescence analysis with a standard multiwell plate reader (Tecan, Safire 2; excitation and emission wavelengths: 630 nm and 660 nm). The fluorescence signals were related to the corresponding MSN mass using standard curves, which were prepared from blank liver and flank tumor tissues of non-exposed mice spiked with known amount of MSN and processed according to the same protocol described above (cage control). The prerequisite for reliable dosimetry is that the homogenization and drying process does not destroy the fluorescence signal of the MSNs. This was proven by comparison of fluorescence signals of homogenates from dried and non-dried samples as well as by adding MSN prior and after homogenization. For analysis of a potential enrichment of MSN_tEGFR_ over MSN_AVI_ in the tumor, the MSN concentration (MSN mass per mass of tissue) was calculated for both tumor and liver samples.

#### Statistical analysis

All values are expressed as means ± standard error of the mean, and statistical analyses were made with GraphPad Prism 8 software (San Diego, USA). Non-parametric Mann-Whitney test was used for flow cytometry quantifications. For multiple comparisons, the parametric test 2-way analysis of variance (ANOVA) was used (see legends to Figures for corresponding comparison tests used). * *P* < 0.05, ** *P* < 0.01, and *** *P* < 0.001, **** *P* < 0.0001 were considered statistically significant.

## Acknowledgements

S.D. and C.M-S. contributed equally to this work. SM, DEW and TB conceived the study, and in assistance with DAB, SD, SL, TS, OS, GTS, SvR designed or co-designed all experiments presented. DAB, DEW, SD, SL, SM and TB wrote the manuscript. SD, DG, and CA synthesized and characterized the nanoparticles. ML, OE provided human tissue for histological analysis. DAB and CM-S performed in vitro and in vivo characterization of the nanoparticles. MV generated and monitored the flank tumor models, applied the particles intravenously. LY and OS performed the fluorescence analysis from flank tumor homogenates. DAB, CM, AD, and DEW designed and performed all experiments associated with ORVD. We thank the Nanosystems Initiative Munich (NIM) for providing funding for DAB. The Knut and Alice Wallenberg foundation is acknowledged for generous support (DEW) as well as the Åke och Inger Bergkvists Stiftelse (DEW). Financial support from the Center for NanoScience Munich (CeNS), the DFG (SFB 749 and SFB 1032), and the European Union’s Horizon 2020 research and innovation program under Grant No. 686098 (SmartNanoTox) and the European Research Council (2010 Starting Independent Investigator and 2015 Proof of Concept Grants, #260524 and #679345, respectively, to GTS) is gratefully acknowledged. We thank David Kutschke, Christina Lukas for their generous help as well as Lina Gefors and Sebastian Wasserstrom at the Lund University Bioimaging Center (LBIC) for their support with electron microscopy. We are grateful to Manfred Ogris for stimulating discussions and Hani Alsafadi for graphical design. We also wish to thank all the other members of both the Meiners and Wagner labs for their support and critical discussion.

## Table of Contents (ToC)

Despite their immense potential, clinical translation of nanomedicines has been hampered by physicochemical and biological barriers impairing cell-specific targeting especially in solid tumors. This study reports a novel approach using organ-restricted vascular delivery (ORVD) with direct administration and recirculation of nanoparticles to enhance nanoparticle uptake in lung tumors. ORVD opens up new avenues for optimized nanotherapies.

## Organ-restricted vascular delivery of nanoparticles for lung cancer therapy

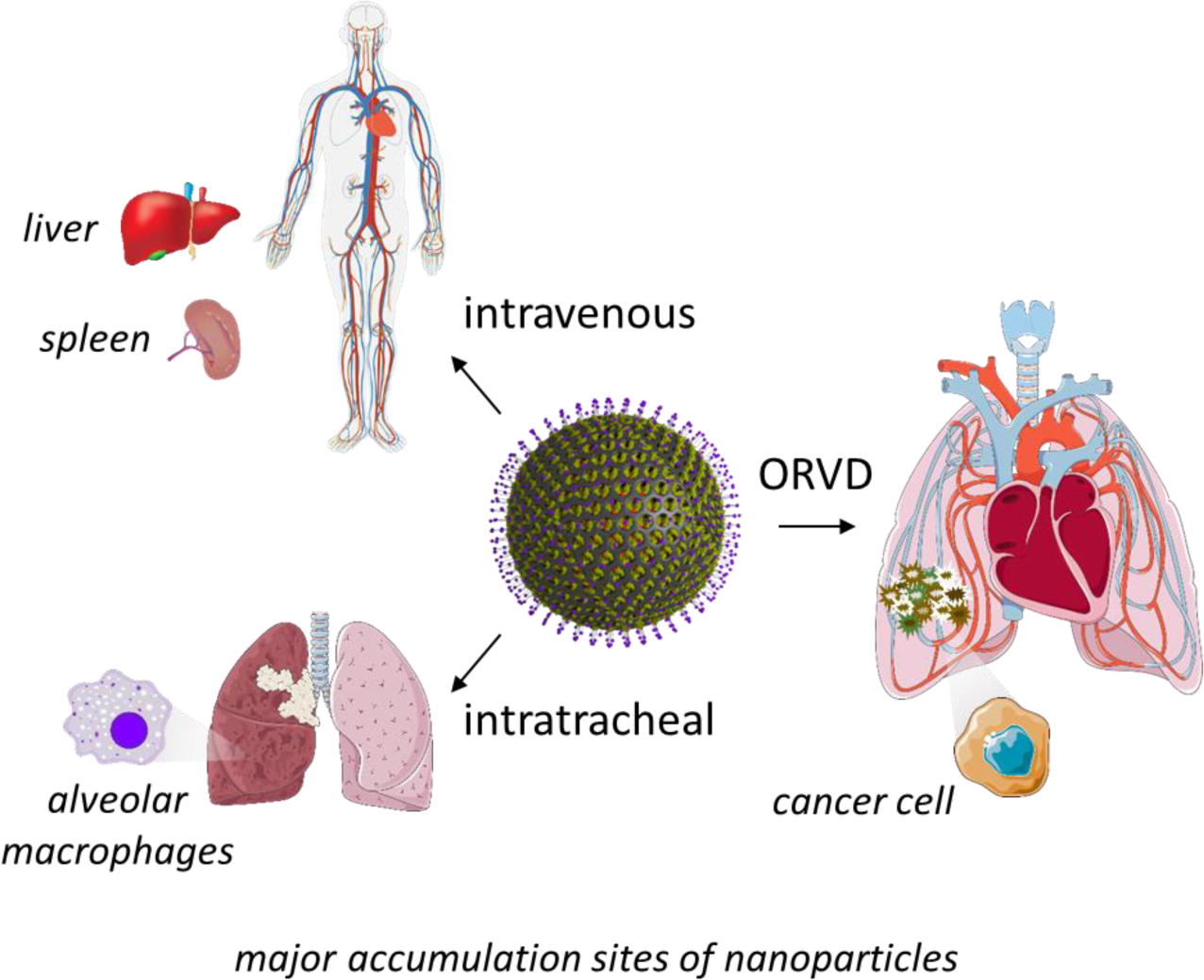

## Supporting Information

**Figure S1.**
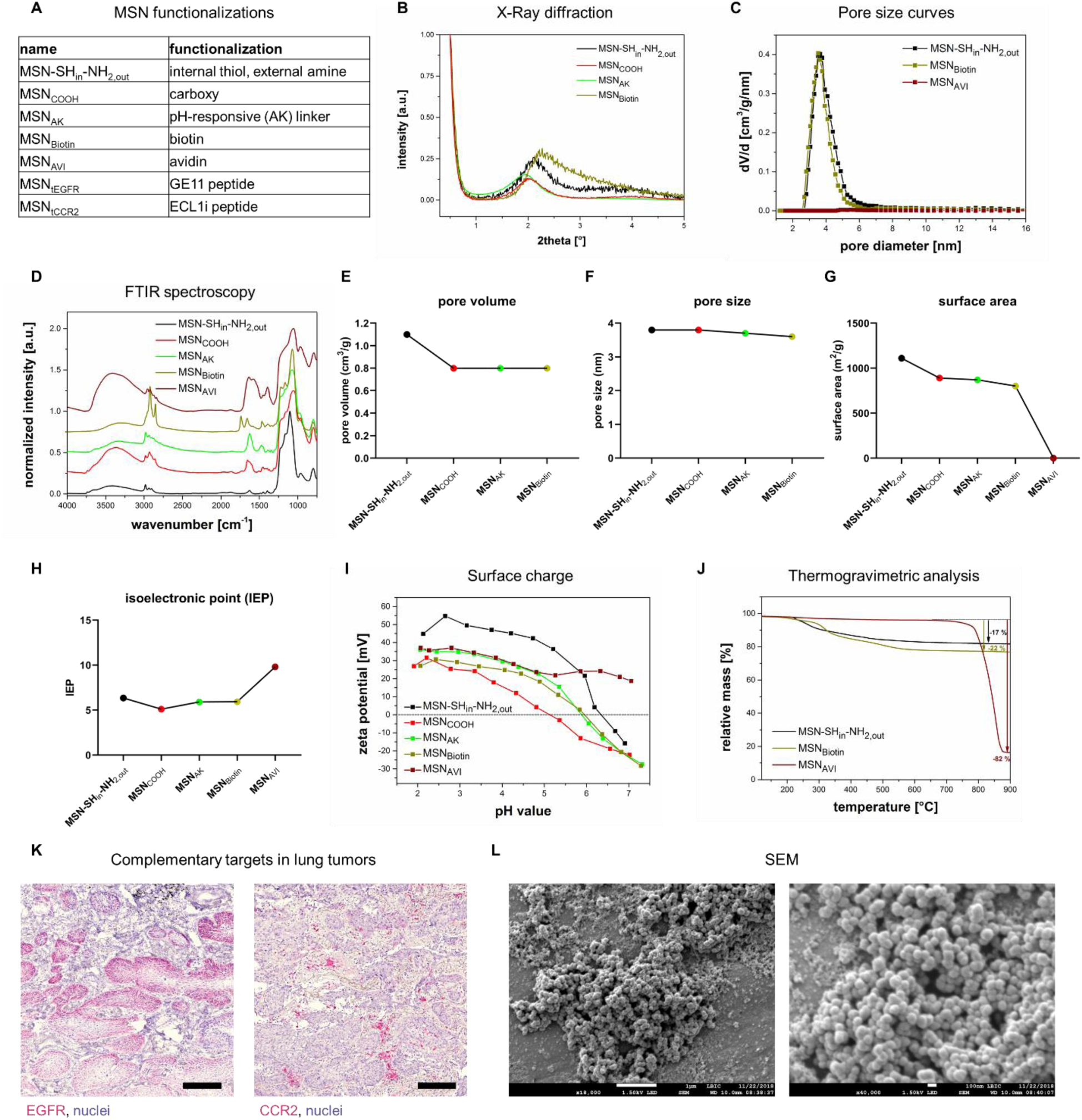
Physicochemical characterization of the MSNs. a) Summary of MSN functionalizations used. b) Small-angle X-ray scattering showing amorphous properties, c) pore diameter distribution curves, d) infrared spectroscopy, e) pore volume, f) pore size, g) surface area, h) isoelectronic point (IEP), i) zeta potential measurements, j) thermogravimetric analysis for the different functionalization stages. k) Immunohistochemical staining representing complementary distribution of EGFR and CCR2 in a human non-small cell lung cancer (NCSLC) specimen. Scale bar = 200 μm. l) Scanning electron microscopic images of MSN_tEGFR_. Scale bar = 1 μm (left) and 100 nm (right).

**Figure S2.**
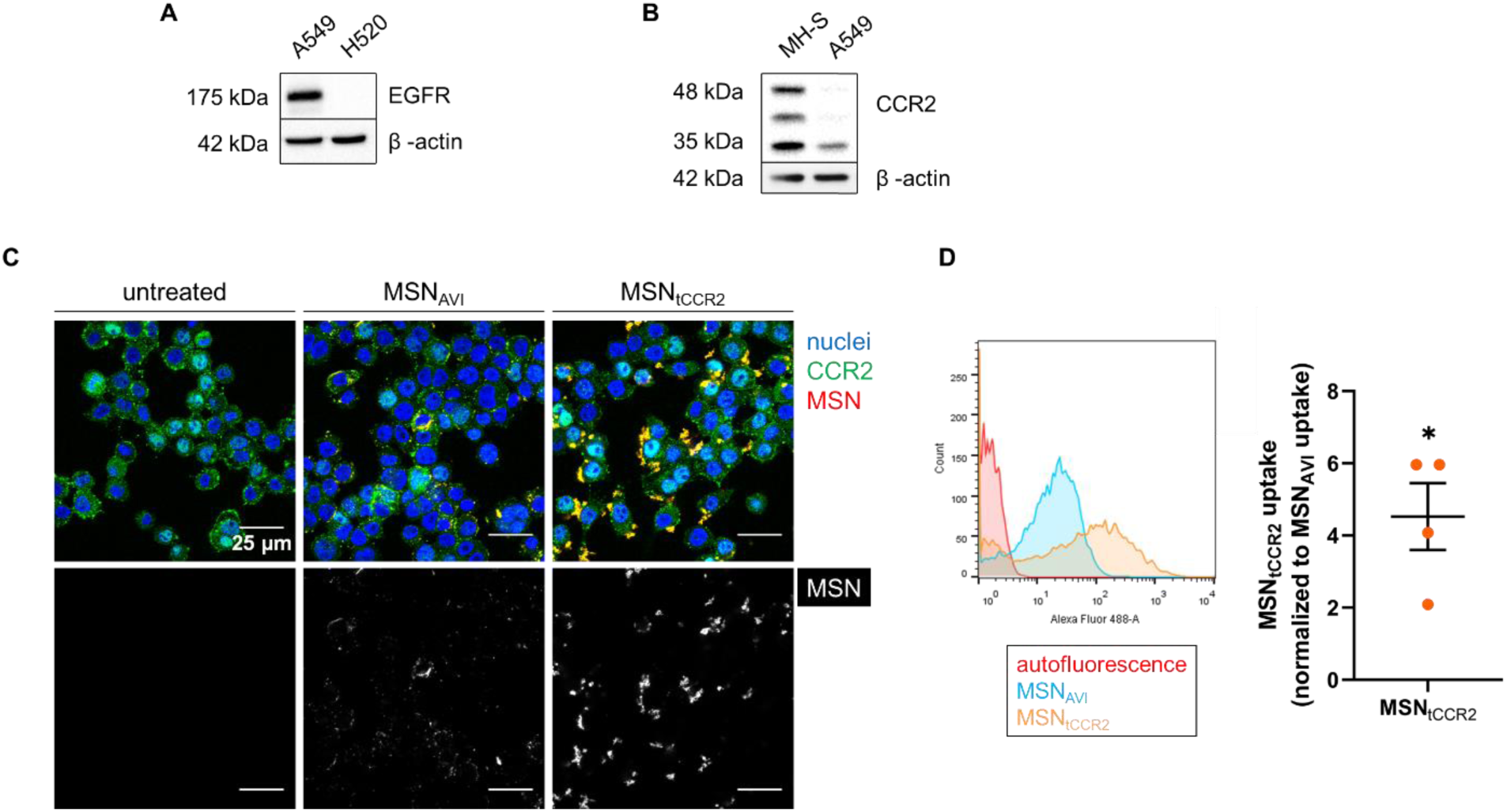
Increased uptake of CCR2 targeted nanoparticles in murine alveolar macrophages *in vitro*. a) EGFR expression in A549 cells in comparison to another NSCLC cell line, H520 cells, b) Western blot analysis of CCR2 expression in murine alveolar macrophage (MH-S) cells in comparison to A549 cells. c) Untargeted *versus* CCR2-targeted uptake of ATTO 633-labeled MSN_AVI_ and MSN_tCCR2_ (red in the upper panel, gray in the lower panel) by MH-S cells after 1 h; CCR2 (green) and cell nuclei (blue) visualized by confocal microscopy. Scale bar = 25 μm. d) Increased uptake of ATTO 488-labeled MSN_tCCR2_ *versus* MSN_AVI_ after 1 h by MH-S cells measured by flow cytometry analysis. * *p* = 0.0286, Mann-Whitney test; values given are an average of four independent experiments ± standard error of the mean.

**Figure S3.**
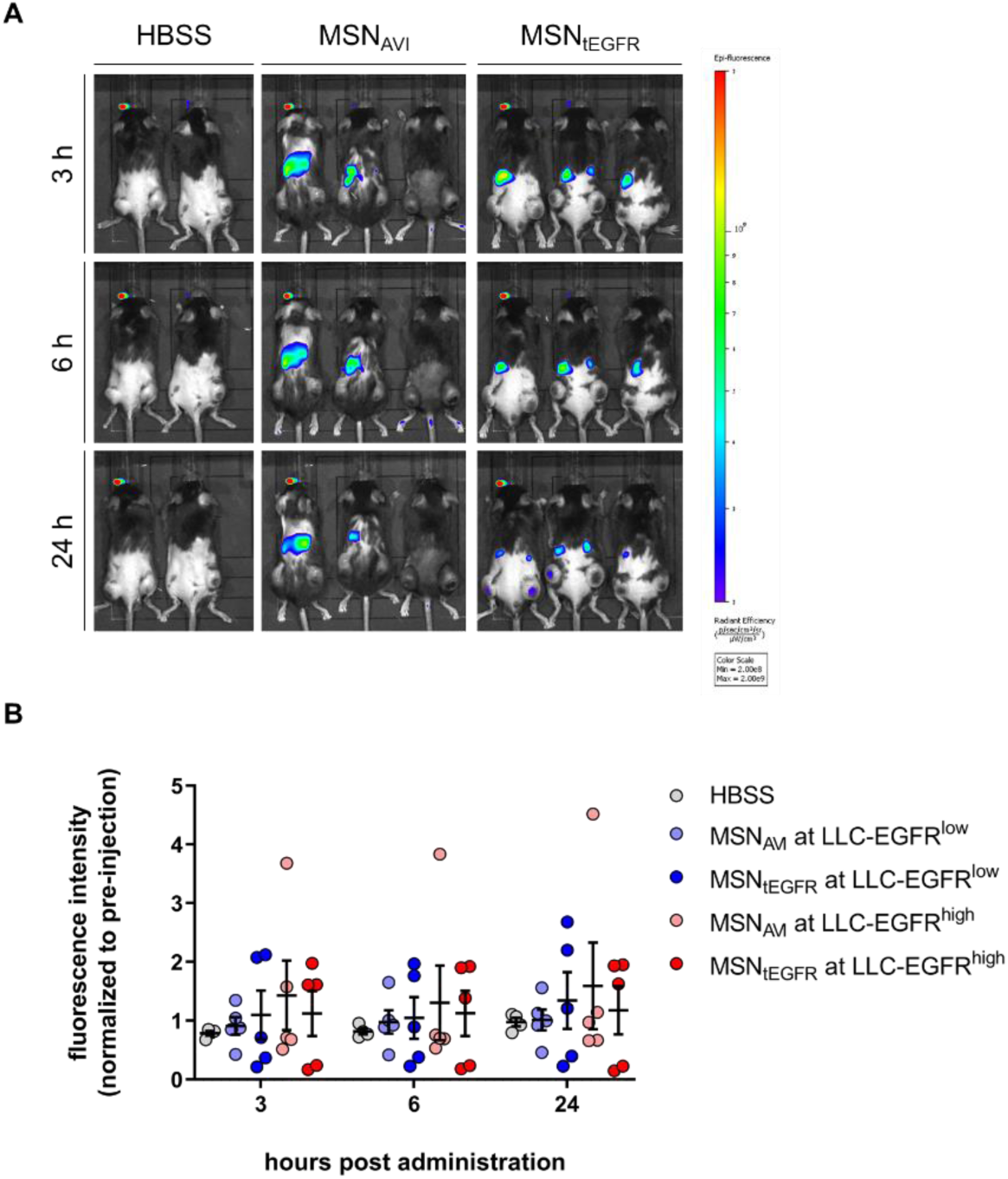
Intravenously administered MSNs are distributed unspecifically in a syngeneic LLC tumor model *in vivo*. a) Representative fluorescence images of mice receiving 1 mg of MSN_AVI_ or MSN_tEGFR_ at 3, 6, and 24 h after retro-orbital administration. b) Quantification of the fluorescence intensity obtained from the individual flank tumors of the mice treated with the MSNs in time course. Values given are an average of signal obtained from five independent mice at each time point ± standard error of the mean (n = 5 mice per MSN type).

**Figure S4.**
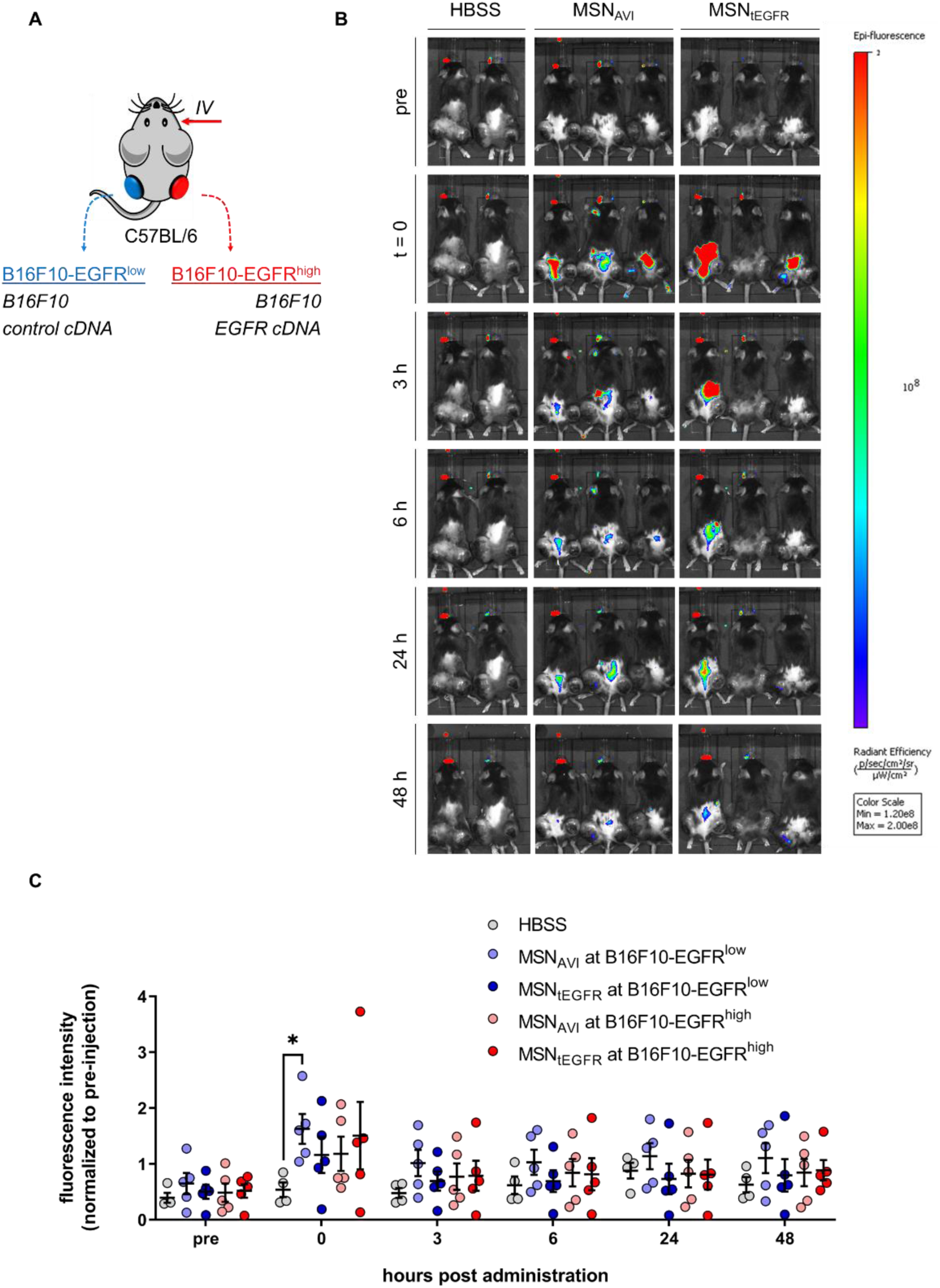
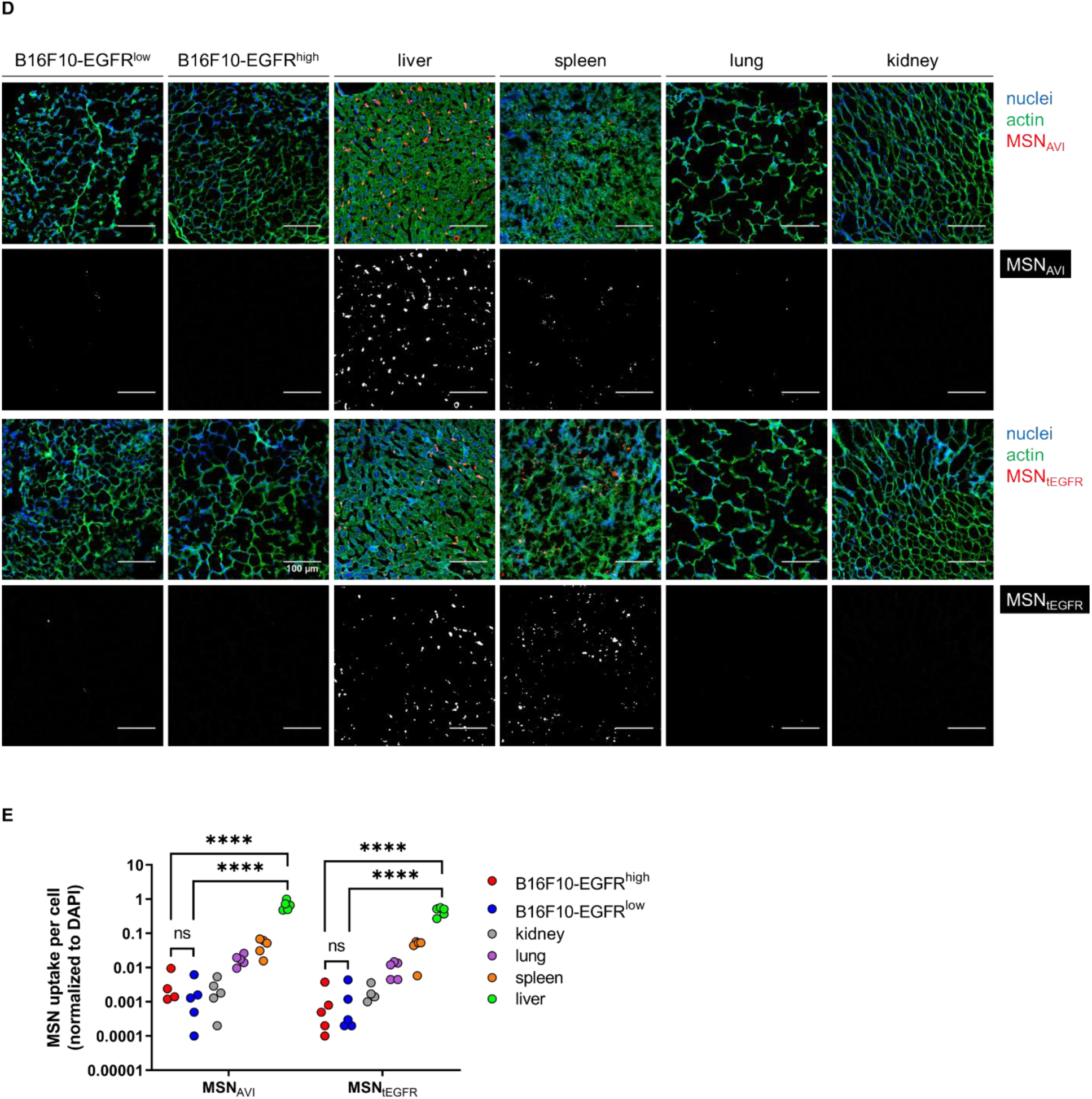
Intravenously administered MSNs are deposited in the liver and spleen in a syngeneic melanoma tumor model *in vivo*. a) Schematic representation of the syngeneic double flank tumor-bearing mouse model that was generated by subcutaneous injection of genetically modified melanoma clones (B16F10) for basal EGFR expression (B16F10-EGFR^low^) *versus* overexpression (B16F10-EGFR^high^). b) Representative fluorescence images of mice receiving 1 mg of MSN_AVI_ or MSN_tEGFR_ before, immediately after, or at 3, 6, 24, and 48 h after retro-orbital administration. c) Quantification of the fluorescence intensity obtained from the individual flank tumors of the mice treated with the MSNs in time course. Values given are an average of signal obtained from five independent mice at each time point ± standard error of the mean. * *p* = 0.0324, Two-way ANOVA, Tukey’s multiple comparisons test. d) Histological analysis for the biodistribution of the intravenously administered MSN_AVI_ and MSN_tEGFR_ in B16F10-EGFR^low^ and B16F10-EGFR^high^ tumors, livers, spleens, lungs, and kidneys of the mice visualized by confocal microscopy. Nuclear staining (DAPI) is shown in blue, cell morphology via actin staining (phalloidin) in green and ATTO 633-labeled MSNs in red in the merged image, and in gray in the single channel. Images shown are representative for three different regions from each mice (n = 5 mice treated). Scale bar = 100 μm. e) Quantification of the MSN_AVI_ and MSN_tEGFR_ uptake per nuclei observed in histological analyses in B16F10-EGFR^low^ and B16F10-EGFR^high^ tumors, kidneys, lungs, spleens, and livers, respectively. In the HBSS control, animals only received HBSS and no particles. Values given are average of three different images per each treated mice ± standard error of the mean (n = 5 per MSN type). **** *p* < 0.0001, Two-way ANOVA, Tukey’s multiple comparisons test.

**Figure S5.**
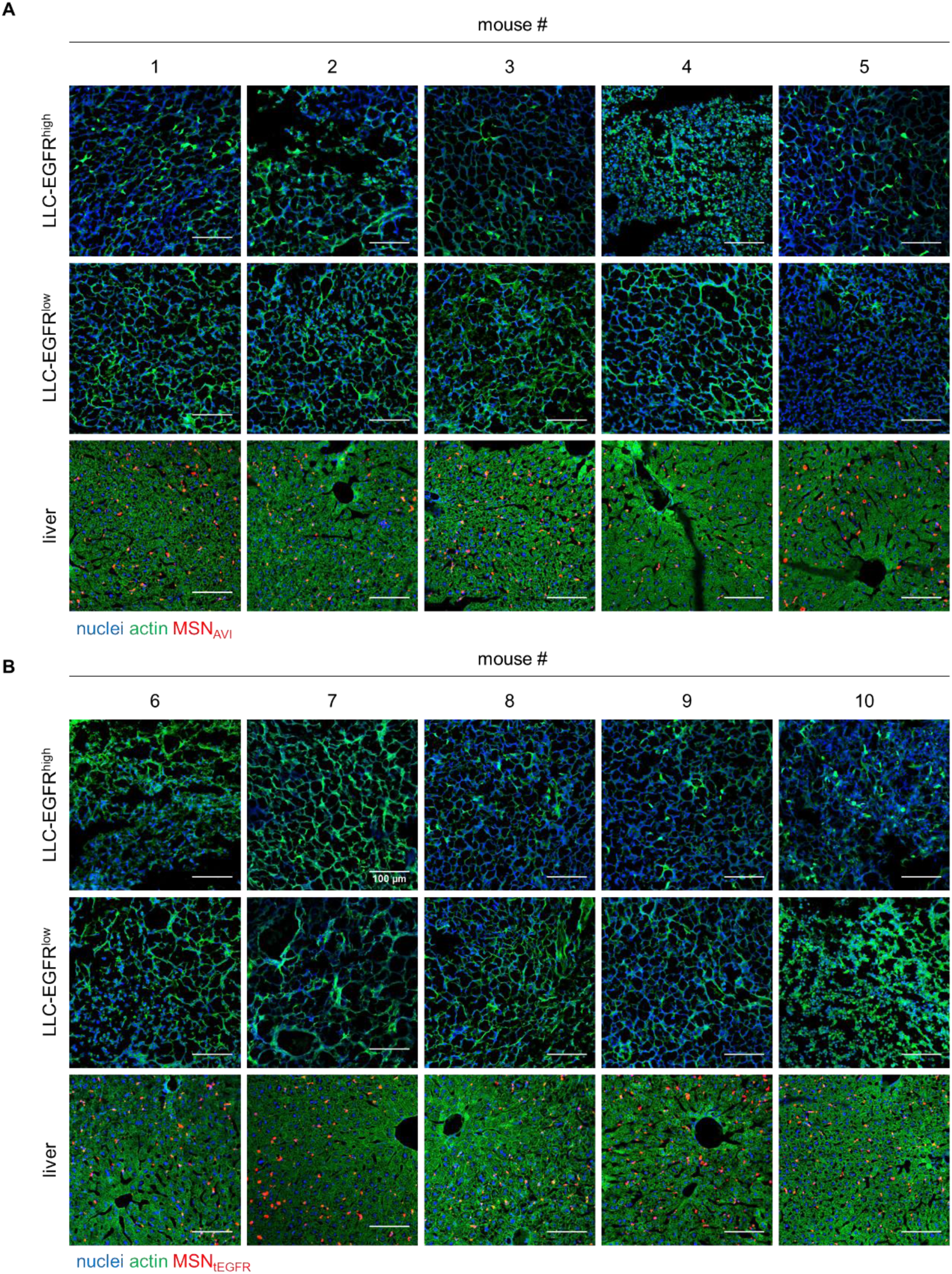
Intravenously administered MSNs are deposited to liver in a syngeneic LLC tumor model *in vivo*. Histological analysis for the biodistribution of the retro-orbitally administered a) MSN_AVI_ and b) MSN_tEGFR_ in the LLC-EGFR^high^ tumors, LLC-EGFR^low^ tumors, and livers of each treated mice by confocal microscopy. Nuclear staining (DAPI) is shown in blue, cell morphology via actin staining (phalloidin) in green, and ATTO 633-labeled MSNs in red. Images shown are representative for three different regions from each mice (n = 5 mice per MSN type). Scale bar = 100 μm.

**Figure S6.**
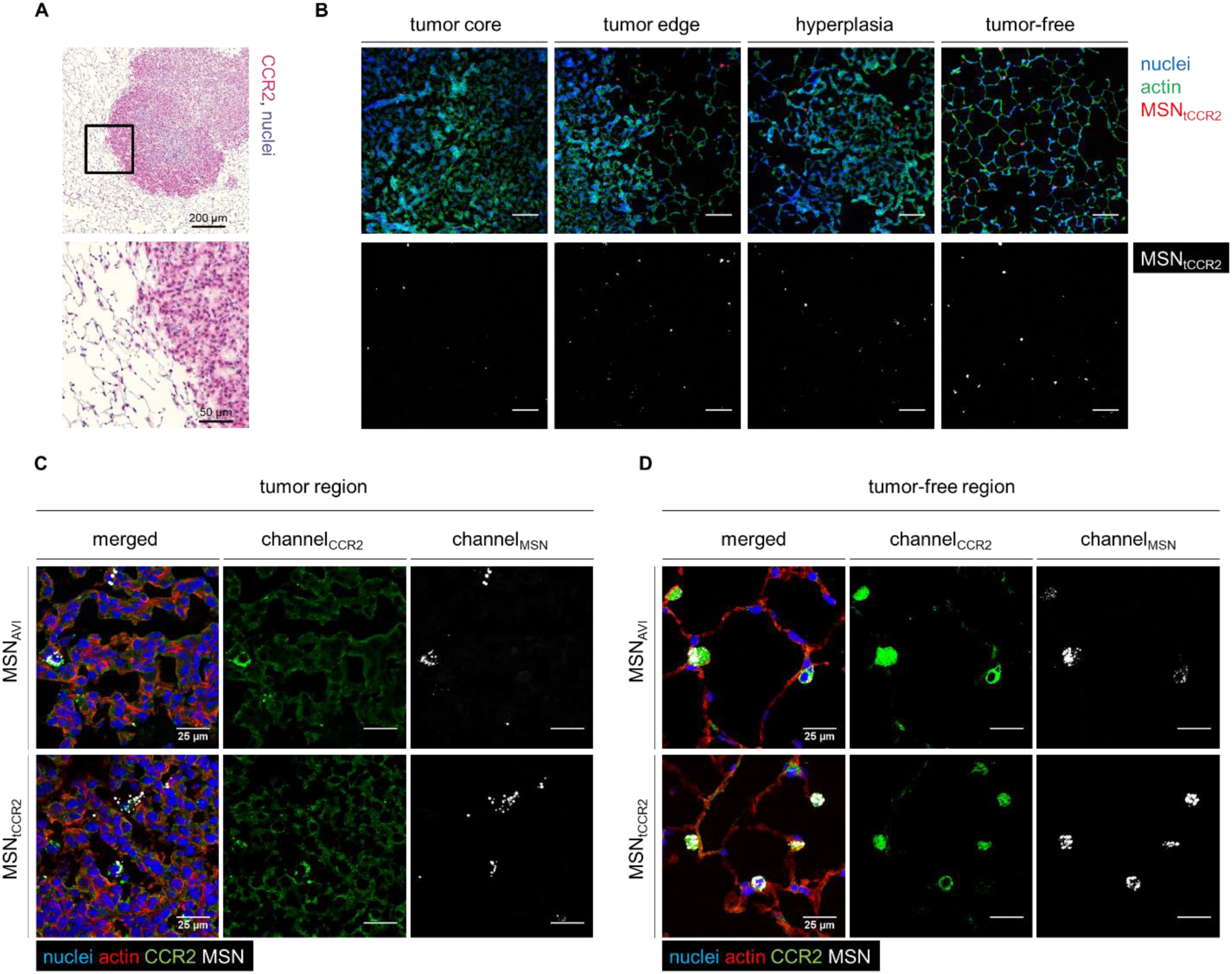
Intratracheally administered MSN_tCCR2_ are engulfed by alveolar macrophages in a mouse model of lung cancer *in vivo*. a) Immunohistochemistry staining of CCR2 (pink) in *Kras^LA2^* mutant tumorous mouse lungs. b) Representative histological analysis of intratracheally instilled ATTO 633-labeled MSN_tCCR2_ uptaken in solid tumor cores *versus* their edges, and in hyperplastic or in tumor-free regions of the tumorous mouse lungs, after 3 days. Nuclear staining (DAPI) is shown in blue, cell morphology via actin staining (phalloidin) in green, and ATTO 633-labeled MSNs in red in the merged images, and in gray in the single channels. Five mice were analyzed per group with five random sections and three images per section in a blinded manner. Images shown are representative for three different regions from each group of mice (n = 5 per MSN type). Scale bar = 100 μm. Immunofluorescence co-staining for CCR2 in c) tumorous *versus* d) tumor-free regions the mutant lungs treated with ATTO 633-labeled MSN_AVI_ *versus* MSN_tCCR2_. Nuclear staining (DAPI) is shown in blue, cell morphology via actin staining (phalloidin) in red, CCR2 staining in green, and ATTO 633-labeled MSNs in gray. Images shown are representative for three different regions from each group of mice (n = 5 per MSN type). Scale bar = 25 μm.

**Figure S7.**
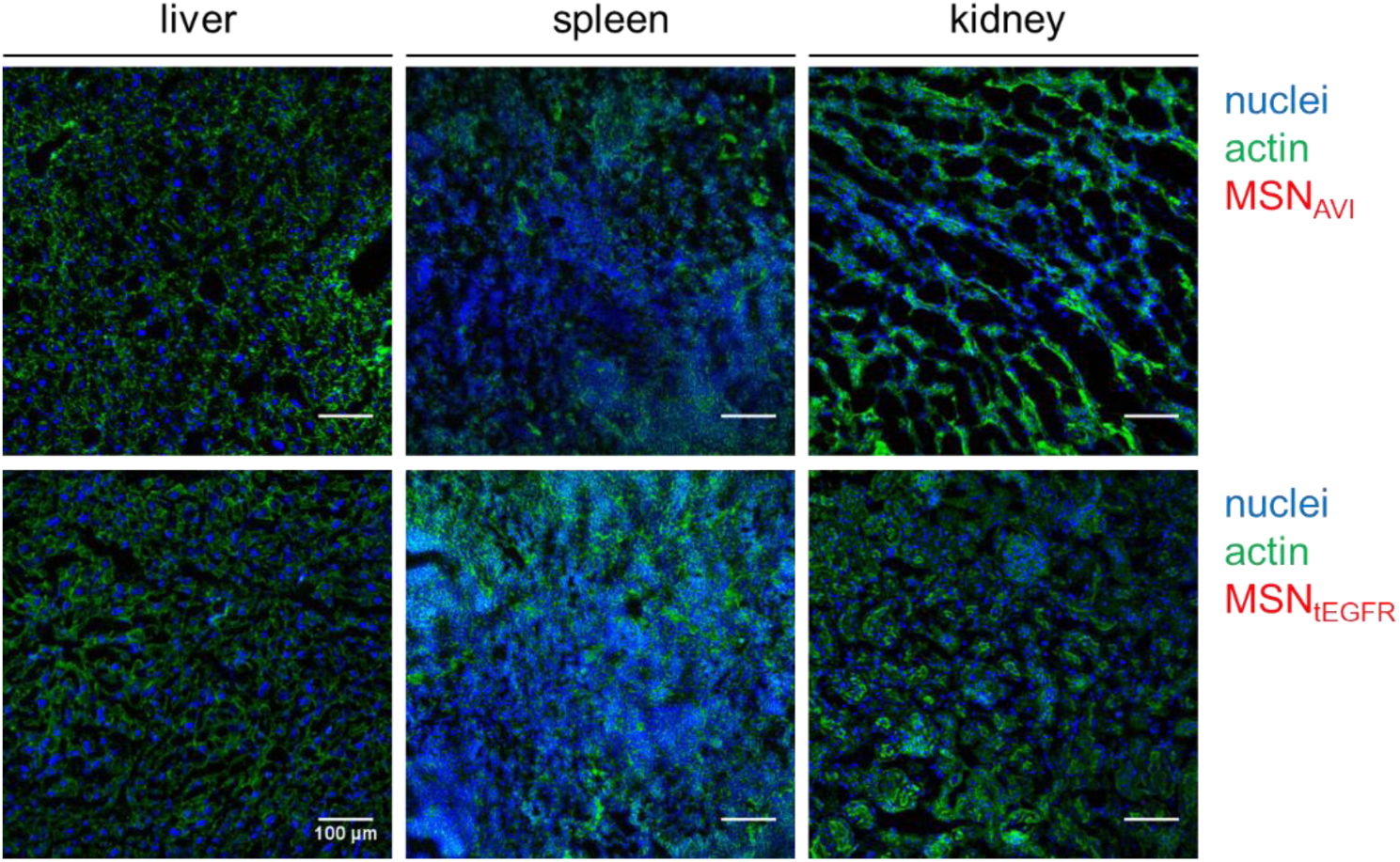
MSN_AVI_ and MSN_tEGFR_ do not localize to the liver, spleen or kidney when administered intratracheally into *Kras^LA2^* mutant mice. Representative histological analysis of intratracheally administered ATTO 633-labeled MSN_AVI_ and MSN_tEGFR_ are not present in livers, spleens, and kidneys of the *Kras^LA2^* mutant mice 3 days. Nuclear staining (DAPI) is shown in blue, cell morphology via actin staining (phalloidin) in green, and ATTO 633-labeled MSNs in red. Images shown are representative for three different regions from each mice (n = 5 per MSN type). Scale bar = 100 μm.

**Figure S8.**
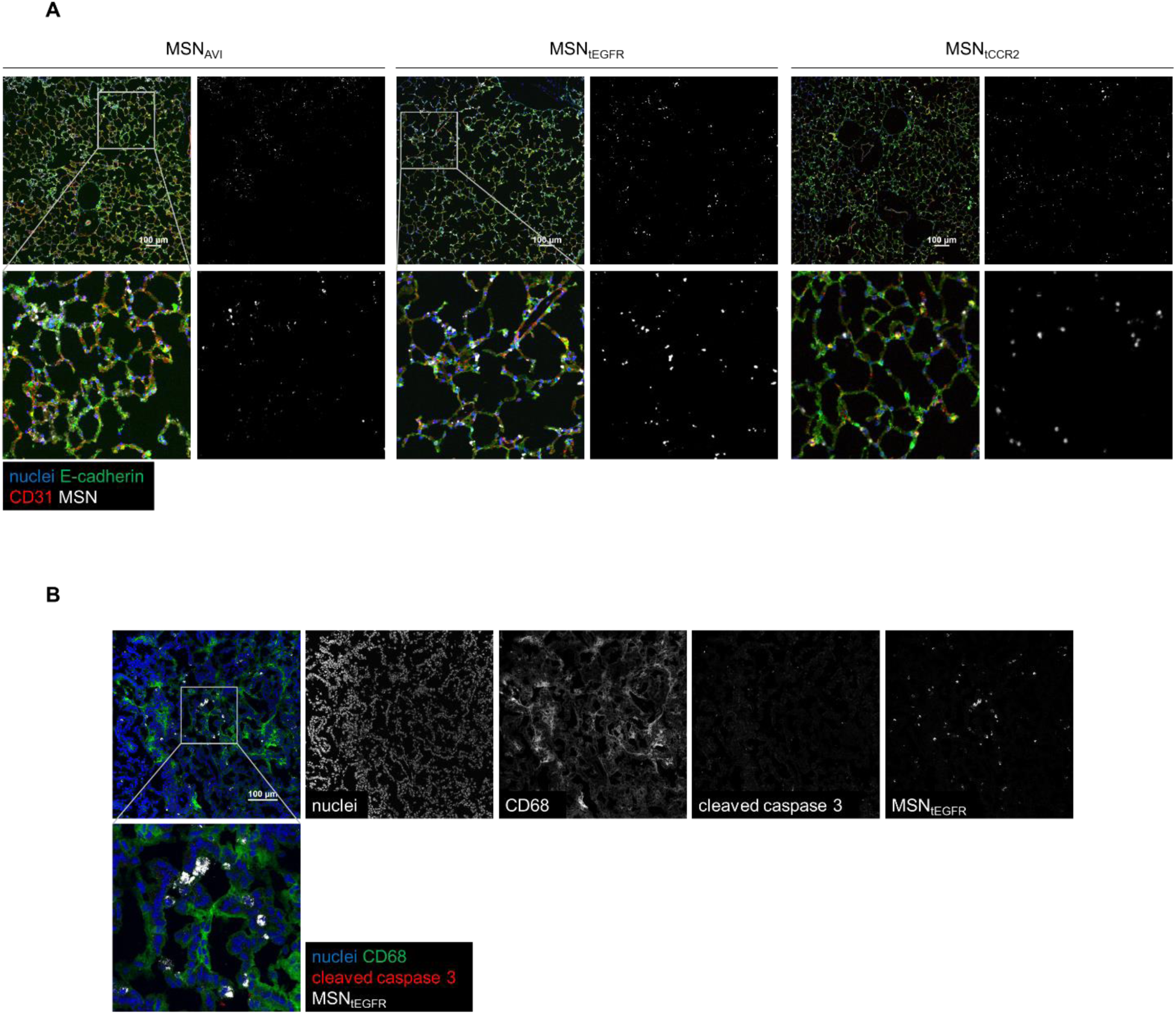
Organ-restricted vascular delivery of MSNs. a) Representative histological analysis of the lungs from WT mice exposed to ATTO 633-labeled MSN_AVI_, MSN_tEGFR_, and MSN_tCCR2_ (all gray). Nuclei were stained with DAPI (blue), epithelial cells labeled with E-Cadherin (green) and endothelial cells labeled with CD31 (red) shown in merged and corresponding single channel images for the nanoparticles. Scale bar = 100 μm. b) Histological analysis of the solid tumors from the *Kras^LA2^* mutant lungs exposed to ATTO 633-labeled MSN_tEGFR_ (gray). Nuclei were stained with DAPI (blue), monocytes/macrophages labeled with CD68 (green) and apoptotic cells labeled with cleaved caspase-3 (red) shown in merged and corresponding single channel images. Scale bar = 100 μm.

## References

[1] J. Fang, H. Nakamura, H. Maeda, Adv. Drug Deliv. Rev. 2011, 63, 136.

[2] D. A. Bolukbas, S. Meiners, Nanomedicine (Lond) 2015, 10, 3203.

[3] D. Rosenblum, N. Joshi, W. Tao, J. M. Karp, D. Peer, Nat. Commun. 2018, 9, 1410.

[4] J.-W. Yoo, D. J. Irvine, D. E. Discher, S. Mitragotri, Nat. Rev. Drug Discov. 2011, 10, 521.

[5] M. J. Ernsting, M. Murakami, A. Roy, S.-D. Li, J. Control. Release 2013, 172, 782.

[6] P. A. Netti, D. A. Berk, M. A. Swartz, A. J. Grodzinsky, R. K. Jain, Cancer Res. 2000, 60, 2497.

[7] K. M. Tsoi, S. A. MacParland, X.-Z. Ma, V. N. Spetzler, J. Echeverri, B. Ouyang, S. M. Fadel, E. A. Sykes, N. Goldaracena, J. M. Kaths, J. B. Conneely, B. A. Alman, M. Selzner, M. A. Ostrowski, O. A. Adeyi, A. Zilman, I. D. McGilvray, W. C. W. Chan, Nat. Mater. 2016, 15, 1212; M. Geiser, J Aerosol Med Pulm Drug Deliv 2010, 23, 207.

[8] S. Wilhelm, A. J. Tavares, Q. Dai, S. Ohta, J. Audet, H. F. Dvorak, W. C. W. Chan, Nat. Rev. Mater. 2016, 1, 16014.

[9] A. J. Tavares, W. Poon, Y.-N. Zhang, Q. Dai, R. Besla, D. Ding, B. Ouyang, A. Li, J. Chen, G. Zheng, C. Robbins, W. C. W. Chan, P. Natl. Acad. Sci. USA 2017, 114, E10871.

[10] F. Bray, J. Ferlay, I. Soerjomataram, R. L. Siegel, L. A. Torre, A. Jemal, CA-Cancer J. Clin. 2018, 68, 394.

[11] D. B. Doroshow, M. F. Sanmamed, K. Hastings, K. Politi, D. L. Rimm, L. Chen, I. Melero, K. A. Schalper, R. S. Herbst, Clin. Cancer Res. 2019, DOI: 10.1158/1078-0432.ccr-18-1538clincanres.1538.2018.

[12] J.-P. Sculier, T. Berghmans, A.-P. Meert, Eur. Respir. Rev. 2015, 24, 23.

[13] P. Zarogoulidis, E. Chatzaki, K. Porpodis, K. Domvri, W. Hohenforst-Schmidt, E. P. Goldberg, N. Karamanos, K. Zarogoulidis, Int J Nanomedicine 2012, 7, 1551.

[14] T. L. Demmy, Video-assist. Thorac. Surg. 2018, 3.

[15] A. Ward, K. Prokrym, H. Pass, Thoracic surgery clinics 2016, 26, 55.

[16] S. Sindhwani, A. M. Syed, J. Ngai, B. R. Kingston, L. Maiorino, J. Rothschild, P. MacMillan, Y. Zhang, N. U. Rajesh, T. Hoang, J. L. Y. Wu, S. Wilhelm, A. Zilman, S. Gadde, A. Sulaiman, B. Ouyang, Z. Lin, L. Wang, M. Egeblad, W. C. W. Chan, Nat. Mater. 2020, DOI: 10.1038/s41563-019-0566-2.

[17] S. H. van Rijt, D. A. Bölükbas, C. Argyo, S. Datz, M. Lindner, O. Eickelberg, M. Königshoff, T. Bein, S. Meiners, ACS nano 2015, 9, 2377.

[18] N. Normanno, A. De Luca, C. Bianco, L. Strizzi, M. Mancino, M. R. Maiello, A. Carotenuto, G. De Feo, F. Caponigro, D. S. Salomon, Gene 2006, 366, 2.

[19] A. Schmall, H. M. Al-Tamari, S. Herold, M. Kampschulte, A. Weigert, A. Wietelmann, N. Vipotnik, F. Grimminger, W. Seeger, S. S. Pullamsetti, R. Savai, Am J Respir Crit Care Med 2015, 191, 437.

[20] Z. Li, R. Zhao, X. Wu, Y. Sun, M. Yao, J. Li, Y. Xu, J. Gu, FASEB j 2005, 19, 1978.

[21] C. Auvynet, C. Baudesson de Chanville, P. Hermand, K. Dorgham, C. Piesse, C. Pouchy, L. Carlier, L. Poupel, S. Barthelemy, V. Felouzis, C. Lacombe, S. Sagan, S. Chemtob, C. Quiniou, B. Salomon, P. Deterre, F. Sennlaub, C. Combadiere, FASEB j 2016, 30, 2370.

[22] T. Agalioti, A. D. Giannou, A. C. Krontira, N. I. Kanellakis, D. Kati, M. Vreka, M. Pepe, M. Spella, I. Lilis, D. E. Zazara, E. Nikolouli, N. Spiropoulou, A. Papadakis, K. Papadia, A. Voulgaridis, V. Harokopos, P. Stamou, S. Meiners, O. Eickelberg, L. A. Snyder, S. G. Antimisiaris, D. Kardamakis, I. Psallidas, A. Marazioti, G. T. Stathopoulos, Nat. Commun. 2017, 8, 15205.

[23] Q. Liu, M. Das, Y. Liu, L. Huang, Adv. Drug Deliv. Rev. 2018, 127, 208.

[24] S. H. van Rijt, T. Bein, S. Meiners, Eur. Respir. J. 2014, 44, 765.

[25] L. Johnson, K. Mercer, D. Greenbaum, R. T. Bronson, D. Crowley, D. A. Tuveson, T. Jacks, Nature 2001, 410, 1111.

[26] D. A. Bölükbas, M. M. De Santis, H. N. Alsafadi, A. Doryab, D. E. Wagner, in Mouse Cell Culture: Methods and Protocols, DOI: 10.1007/978-1-4939-9086-3_20 (Ed: I. Bertoncello), Springer New York, New York, NY 2019, p. 275.

[27] N. M. Weathington, D. Álvarez, J. Sembrat, J. Radder, N. Cárdenes, K. Noda, Q. Gong, H. Wong, J. Kolls, J. D’Cunha, R. K. Mallampalli, B. B. Chen, M. Rojas, JCI Insight 2018, 3, e95515.

[28] R. S. Herbst, D. Morgensztern, C. Boshoff, Nature 2018, 553, 446.

[29] D. F. Quail, J. A. Joyce, Nat Med 2013, 19, 1423.

[30] L. Lang-Lazdunski, Eur. Respir. Rev. 2013, 22, 382.

[31] H. Uramoto, F. Tanaka, Transl. Lung. Cancer Res. 2014, 3, 242.

[32] J. Cosaert, E. Quoix, Br. J. Cancer 2002, 87, 825.

[33] R. K. Jain, T. Stylianopoulos, Nat Rev Clin Oncol 2010, 7, 653.

[34] S. H. van Rijt, D. A. Bölükbas, C. Argyo, K. Wipplinger, M. Naureen, S. Datz, O. Eickelberg, S. Meiners, T. Bein, O. Schmid, T. Stoeger, Nanoscale 2016, 8, 8058.

[35] K. C. Valkenburg, A. E. de Groot, K. J. Pienta, Nat Rev Clin Oncol 2018, 15, 366.

[36] R. A. de Mello, D. S. Marques, R. Medeiros, A. M. Araujo, World J Clin Oncol 2011, 2, 367.

[37] C. Argyo, V. Weiss, C. Bräuchle, T. Bein, Chem. Mater. 2014, 26, 435.

[38] I. Ferrer, J. Zugazagoitia, S. Herbertz, W. John, L. Paz-Ares, G. Schmid-Bindert, Lung Cancer 2018, 124, 53.

[39] X. Liu, A. Situ, Y. Kang, K. R. Villabroza, Y. Liao, C. H. Chang, T. Donahue, A. E. Nel, H. Meng, ACS nano 2016, 10, 2702; M. E. Davis, J. E. Zuckerman, C. H. Choi, D. Seligson, A. Tolcher, C. A. Alabi, Y. Yen, J. D. Heidel, A. Ribas, Nature 2010, 464, 1067.

[40] J. D. O’Neill, B. A. Guenthart, J. Kim, S. Chicotka, D. Queen, K. Fung, C. Marboe, A. Romanov, S. X. L. Huang, Y.-W. Chen, H.-W. Snoeck, M. Bacchetta, G. Vunjak-Novakovic, Nat. Biomed. Eng. 2017, 1, 0037.

